# Spatial population structure can reverse the dependence of mutant burden on the death-to-birth ratio of cells

**DOI:** 10.1101/2025.11.17.688895

**Authors:** Samrat S. Mondal, Natalia L. Komarova, Dominik Wodarz

## Abstract

Expanding cell populations, including bacterial colonies and tumors, continuously accumulate mutations as they grow. Yet how mutant burden depends on the cellular death-to-birth ratio in spatially structured populations has not been addressed, despite clinical studies associating higher tumor turnover with both better and worse patient outcomes across cancers. This question is especially relevant for deleterious mutants, which can become advantageous if the environment changes, such as when treatment begins. Mutant accumulation has been studied extensively in well-mixed populations. Here, we derive a new closed-form approximation for the mean deleterious mutant burden at a specified population size when cell death is nonzero, serving as a baseline for comparisons to spatial structure. We show that spatial structure, a defining feature of solid tumors, can fundamentally alter how mutant burden depends on the death-to-birth ratio. Relationships that increase monotonically in well-mixed populations can become nonmonotonic or even decrease in spatially expanding populations, depending on population size and the fitness cost of mutants. These results provide a mechanistic framework for understanding why higher tumor turnover has been associated with either improved or worsened outcomes across patients and cancer types. An analysis of the probability that a colony regrows after an environmental change (such as treatment initiation) reveals a similar pattern: spatial structure can reverse or reshape this dependence. Overall, our findings establish spatial structure as a key determinant of how cellular death-to-birth ratios shape mutant accumulation, with implications for interpreting clinical data and predicting the burden of preexisting mutants during tumor evolution.

**Significance statement:** The number of mutants present when a growing cell population reaches a given size shapes its capacity to adapt, since deleterious mutants can become advantageous once conditions change, such as when cancer treatment begins. Clinical studies report contradictory links between tumor cell turnover (death-to-birth ratio) and patient outcomes, with higher turnover linked to both better and worse prognosis. Building on established theory for well-mixed populations, we show that spatial population structure, a key feature of solid tumors, can make mutant burden increase, decrease, or vary non-monotonically with the death-to-birth ratio, depending on tumor size and mutant fitness cost, patterns absent in the well-mixed system. This resolves the contradiction and clarifies how tumor turnover shapes its potential to evolve and progress.

## 1 Introduction

The accumulation of mutations in expanding cell populations is a fundamental process that determines their potential to adapt. The number of mutant cells when an expanding colony has reached a given size determines the ability of the population to adapt further and overcome selective barriers [1]. In a biomedical context, this is very important for tumors, which have to continuously evolve as they grow for the disease to progress. The size of a mutant clone in a growing colony of cells can correlate with the chances of single-hit mutant cells to accumulate further mutations in the process of multi-stage carcinogenesis. Similarly, the size of drug-resistant mutant clones when therapy is initiated determines how long it takes for the resistant cell colony to grow to detectable levels. If the number of resistant mutant clones is relatively small when therapy is initiated and the survival of the colony is not guaranteed, the probability of mutant (and thus cell colony) survival becomes an important measure, a concept that has been termed evolutionary rescue among evolutionary biologists [2–6].

The distribution of mutant cell numbers in an exponentially growing population was first characterized by [7], whose fluctuation test established that mutations arise spontaneously before selection acts. The resulting heavy-tailed Luria–Delbrück distribution was refined by [8] using a pure-birth stochastic process for clone growth, and the mathematical theory was extended and reviewed over subsequent decades [9–13]. These classical analyses assumed neutral mutants growing at the same rate as the wild type and, crucially, neglected cell death. Extensions incorporating a birth–death process for mutant clones were developed by [14] and [15], the latter allowing mutants with a selective advantage relative to wild-type cells. Using a probability generating-function approach, [16] analyzed the emergence of resistance mutations during exponential tumor growth under a birth–death model, assuming advantageous, neutral, and deleterious mutants, and showed that a higher cell death rate increases both the probability of resistance and the expected number of resistant cells at a given population size, because more cell divisions are required to reach that size. The same conclusion was reached by [17], who studied neutral mutants and derived analytical results for the mean mutant burden as a function of the death-to-birth ratio, and further showed that mutant burden measured at a given time is different than the mutant burden at a specified population size [17, 18]. Only recently, a more comprehensive analysis of fully-stochastic colony growth in the presence of cell death was performed. One of the most relevant contributions was made by [19], who derived an expression for the probability generating function for the mutant number probability distribution, in the presence of cell death. The authors explicitly explored the long-time limit in specific cases, such as advantageous mutants in a growing colony, and found that the mutant numbers had a fat-tailed distribution. In a subsequent study, [20] extended the framework to accommodate outcomes corresponding to a finite system size (but in the absence of death), and also derived a more general long-term (or large-size) limit, conditioned on the absence of jackpot events. Later, [21] considered a generalization to a multitype branching process to show the abundance distribution and arrival time of mutants, again in the large-time limit. Other work that has also examined related topics is mutation accumulation experiments, where repeated bottlenecks are used to minimize selection but mutant lineages nevertheless expand during colony growth between transfers [22, 23].

While this mathematical literature about mutant accumulation in birth–death processes has made significant progress, we so far do not have a comprehensive understanding of how the death-to-birth ratio of cells influences the number of mutants at a given size threshold in an expanding population, depending on assumptions about mutant fitness, population size, and population structure. Yet, this is a biologically important question, especially when considering the evolutionary dynamics of tumors. Cells from different cancers can vary vastly in their death rates relative to their division rates. Pancreatic ductal adenocarcinoma exhibits a relatively high death-to-birth ratio, with substantial cell death balanced by continuous proliferation [24], while other cancers, such as chronic lymphocytic leukemia, have been reported to display lower death-to-birth ratios due to disrupted apoptosis and extended survival of the tumor cells [25]. Within specific malignancies, the balance between apoptosis and proliferation can vary, and this variation has been shown to correlate with clinical aggressiveness in complex ways, with both higher and lower death-to-birth ratios linked to poorer outcomes in different contexts [26–28]. Elucidating how the death-to-birth ratio shapes mutant burden in growing populations is therefore crucial for interpreting such observations, for understanding the evolutionary potential of specific tumors, and ultimately for predicting mutant burden upon treatment initiation and the likely duration of tumor control based on patient-specific parameters.

To fill this knowledge gap, we take the following multi-step approach. We first consider stochastic, exponentially growing populations and present a comprehensive analysis of the relationship between mutant numbers and cell death-to-birth ratios for deleterious as well as advantageous mutants. This analysis paints a more complicated picture than previously described in the literature, and hence substantially extends our understanding of these evolutionary dynamics. We then extend our modeling approaches to spatially structured populations and show how the dependence of mutant numbers on the cellular death-to-birth ratio is altered, compared to well-mixed colonies. In these analyses, we focus mostly on the average number of mutants in a growing cell colony as a measure of interest, because this number has important biological implications, as reviewed above. We finish our analysis by presenting selected results that demonstrate how cell death-to-birth ratio can influence the probability of evolutionary rescue, both in well-mixed and in spatially explicit models. This is relevant for scenarios where mutant numbers are relatively low, such that both population extinction and rescue are likely outcomes.

## 2 Results

We consider a number of models where a population of individuals undergoes unlimited growth through births and deaths, and mutants are generated with a certain probability during cell divisions. Let us denote by *r*_w_ and *d*_w_ the basic, per capita division and death rates of the wild-type cells. The quantity

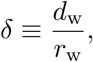

that is, the death-to-birth ratio of the wild-type cells, is at the focus of the current study. We have, *d*_w_ = *δr*_w_. The mutant kinetic parameters are given by *r*_m_ = (1 + *s*_1_)*r*_w_ and *d*_m_ = (1 − *s*_2_)*d*_w_, such that positive values of the selection factors *s*_1_ and *s*_2_ generally align with advantageous, and negative values with disadvantageous mutants.

In the following, we explore how the cellular death-to-birth ratio influences the number of mutants when the growing colony reaches a threshold size. We first describe this relationship a in well-mixed model as a baseline, and then explore how spatial population structure alters the results.

### Death-to-birth ratio and mutant accumulation in a well-mixed colony

Consider a linear birth-death process with mutations, where *x*(*t*) and *y*(*t*) measure the populations of wild-type and mutant cells respectively, and *µ* denotes the mutation rate:

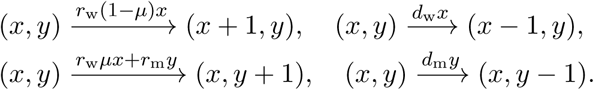

Stochastic simulations of this system were performed as follows. Under the initial condition *x*(0) *>* 0*, y*(0) = 0, we let the population grow to the total size *N*, and evaluate the average mutant burden, ⟨*M_N_* ⟩, as a function of parameters, based on repeated realizations of the simulation. In the following we describe how this mutant burden ⟨*M_N_* ⟩ depends on the death-to-birth ratio, *δ*. To our knowledge, such an analysis has so far not been explicitly described in the literature, even though [20] derived a formula for mutant burden at a (large) threshold size in a stochastic model of exponential population growth in the presence of both birth and death. This analysis was conditioned on the wild-type population non-extinction, which excluded important jackpot mutation events (see below), and it was shown that in this case the results were identical to those given by the corresponding deterministic system, see SI. Equation (1). We will refer to the deterministic prediction for the number of mutants at size *N* as 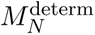.

Here we extend this work by approximating the number of mutants at size *N* in a stochastic model of exponential population growth, taking into account what we call “extreme jackpot events”, i.e. the possibility of an extreme event where the final state is (*X, Y*) = (0*, N*). In general, jackpot events are realizations contributing to the fat tail of the clone-size or mutant-number distribution. Such events occur in well-mixed systems, and they play an even bigger role in spatially growing colonies [29]. Unlike spatial jackpot events, which are typically driven by sector formation, the jackpot events in well-mixed populations arise from rare events when a mutant is generated early on in colony growth. The most extreme of jackpot events is a stochastic extinction of the wild-type lineage, followed by an expansion of a mutant lineage generated prior to extinction. Figure 1(a) shows a numerical approximation to the distribution of the mutant burden at size *N* in a well-mixed population. One can see a nonzero contribution of populations that consist entirely of mutants. This is a signature of the extreme jackpot events.

**Figure 1:**
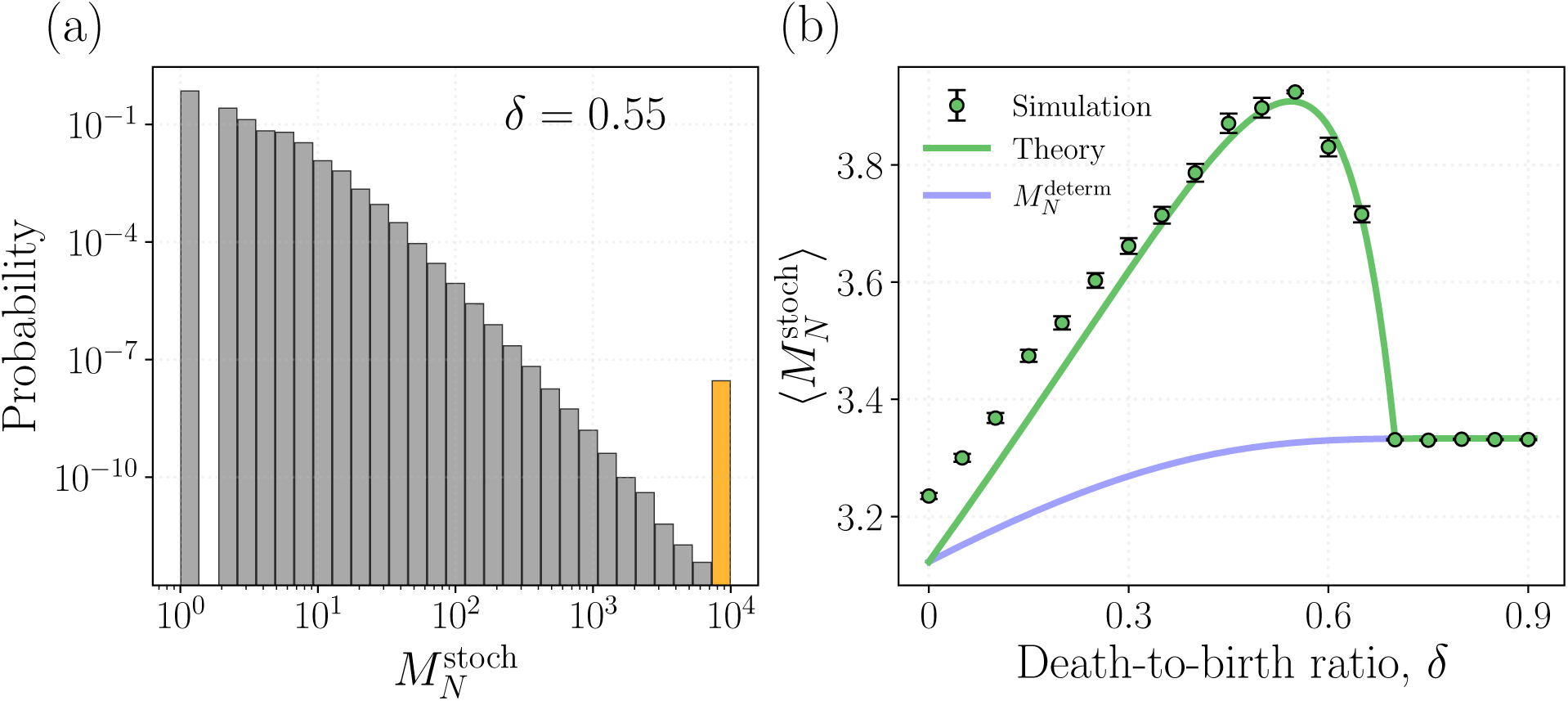
Distribution of mutant burden and dependence of the mean mutant burden on the death-to-birth ratio in a well-mixed stochastic system. Panel (a) depicts the distribution of the final mutant burden *M_N_* at the death-to-birth ratio *δ* = 0.55, obtained from 10^9^ non-extinct runs. The orange-colored bin represents the probability of extreme jackpot events, in which the system consists entirely of mutants. Panel (b) shows the mean mutant burden at different death-to-birth ratios. The blue line represents the deterministic model prediction, the green symbols with error bars (standard error) are stochastic simulations, and the green line is a theoretical approximation, see SI.2.1 for details. For *δ <* (1 + *s*_1_), mutants are supercritical and the mutant burden exceeds the deterministic prediction, whereas for *δ >* (1 + *s*_1_), mutants become subcritical and the burden declines toward the deterministic limit. Each point is computed from 2 × 10^7^ non-extinct runs. For both panels, the other parameters are fixed at *r*_w_ = 0.5, *µ* = 10*^−^*^4^, *N* = 10^4^, and (*s*_1_*, s*_2_) = (−0.3, 0.0).

We theoretically compute (see SI.2.1) the probability of such extreme jackpot events in the limit of rare mutation as follows:

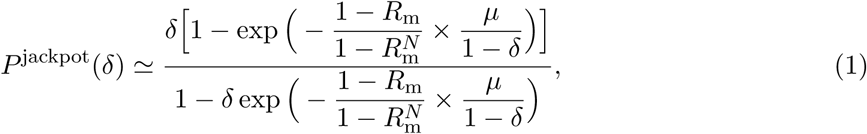

where *R*_m_ = [*δ*(1 − *s*_2_)] */* (1 + *s*_1_). Using this, we obtain a corrected stochastic mutant burden, 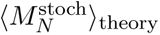:

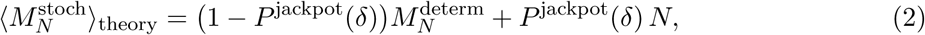

which generally deviates from the deterministic well-mixed prediction *M* ^determ^ (see the green solid line in Figure 1(b)). The mutant burden exceeds the deterministic prediction for lower *δ* due to takeovers of mutants with *r*_m_ *> d*_m_ (supercritical mutants, able to sustain their own growth), but as the death-to-birth ratio increases further, such that *r*_m_ *< d*_m_ (subcritical mutants, not able to sustain their own growth), the mutant burden decreases to coincide with the deterministic prediction, since mutant takeovers become nearly impossible.

We now use this calculation to describe how the death-to-birth ratio affects mutant burden. First, we note that advantageous mutant burden always increases with the death-to-birth ratio (see SI.1.2). The patterns are, however, more complex for deleterious mutants, as illustrated in Figure 2. In this respect, we distinguish between two scenarios: We assume that the mutant disadvantage is expressed in (i) the birth (division) rate of cells (*s*_1_ *<* 0*, s*_2_ = 0, panels (a) and (b)), or (ii) in a difference in the cell death rates (*s*_1_ = 0*, s*_2_ *<* 0, panels (c) and (d)).

**Figure 2:**
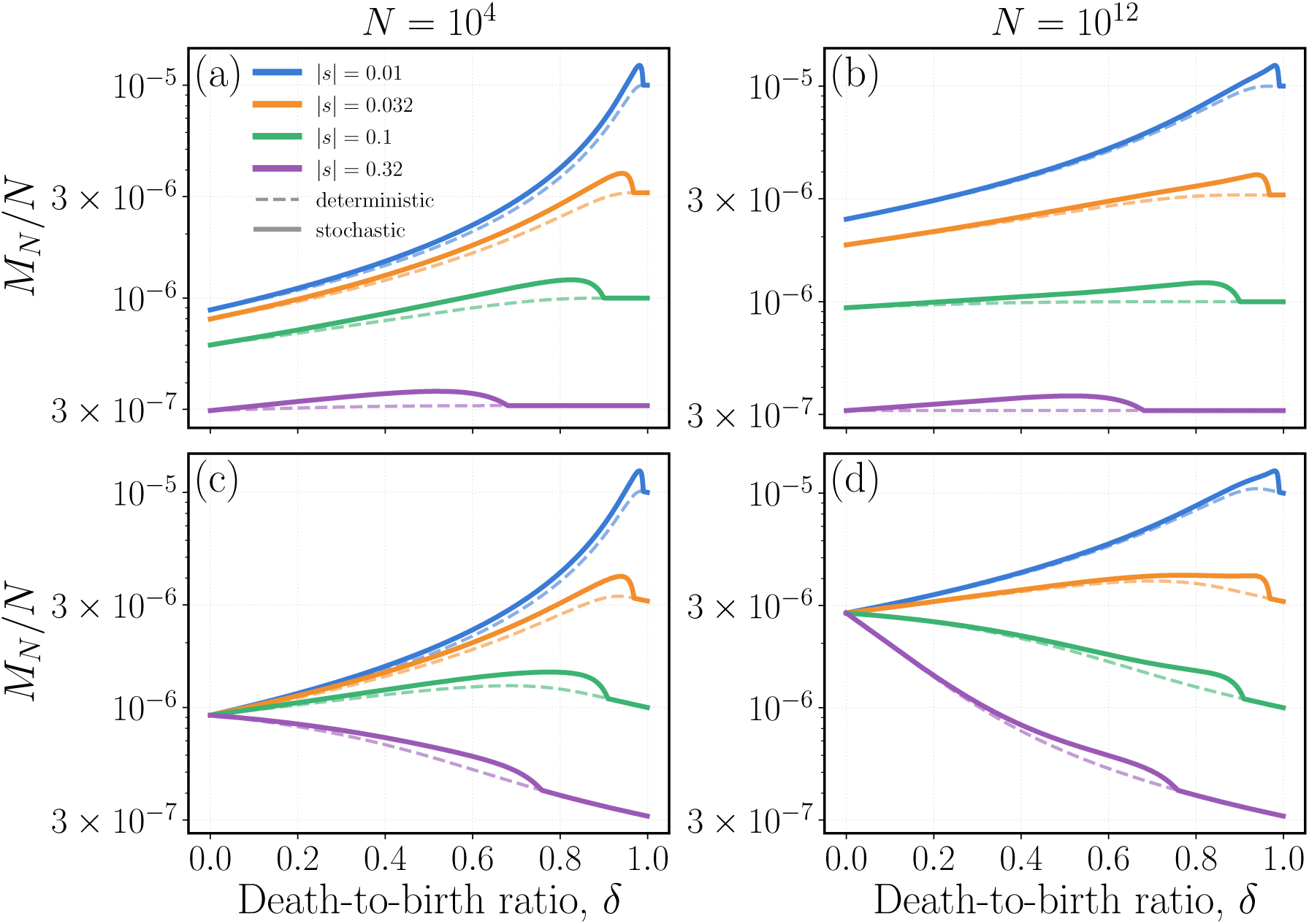
Dependence of mutant burden on the death-to-birth ratio in well mixed systems. Here we compare the ratio of mutant burden to the final population size (*M_N_ /N*) by varying death-to-birth ratio *δ*. The left column (a, c) shows the outcomes for a population size *N* = 10^4^, while the right column (b, d) shows the same outcomes for a final size *N* = 10^12^. The top row (a, b) corresponds to the case where mutants are disadvantageous in birth only, i.e., *s*_2_ = 0, while the bottom row (c, d) represents the case where mutants are disadvantageous in death only, i.e., *s*_1_ = 0. In each panel, four different values of the disadvantage strength are shown in four different colors. The dashed curves represent results from the deterministic model, whereas the solid curves correspond to the stochastic model.

First we focus on a disadvantage in birth. Here we find that over large ranges of *δ* (in which the mutants can sustain their own growth), the number of mutants increases with the death-to-birth ratio, see Figure 2(a,b). The figure shows this relationship for two threshold population sizes and different levels of selective disadvantage. For the lower threshold population size (panel (a)), the number of mutants rises with *δ* for all selection coefficients. Only for very high values of *δ* does this trend reverse, when the mutants transition to becoming unable to sustain their own growth due to high death rates, as explained above and in Figure 1(b). This reversal does not occur in the corresponding deterministic system (dashed lines in Figure 2) due to the lack of jackpot mutation events [20]. For larger threshold population sizes, the relationship between the death-to-birth ratio and mutant burden becomes flatter, although the general trends remain similar (Figure 2(b)).

Next, we assume that the mutant is disadvantageous by having an increased death rate compared to the wild-type (Figure 2(c,d)). This is a more complex scenario because now the selection coefficient *s*_2_ multiplies the death-to-birth ratio *δ*, i.e. the mutant death rate is given by *d*_m_ = (1 + *s*_2_)*δr*_w_. In other words, the higher the death-to-birth ratio, the more pronounced the mutant disadvantage becomes. Consequently, we observe the following patterns: Focusing on lower *δ* ranges (mutants can sustain their own growth), the number of mutants can increase with the death-to-birth ratio, decrease, or show a one-humped relationship depending on the threshold population size and the selective disadvantage of the mutant. For lower threshold population sizes (panel (c)), the mutant burden largely rises with the death-to-birth ratio if the selective disadvantage is relatively low. Larger selective disadvantages can lead to a decrease of mutant numbers with the death-to-birth ratio. For higher threshold population sizes (panel (d)) we see a more pronounced negative relationship between mutant numbers and the death-to-birth ratio over a larger range of selective disadvantages. The reason is that the monotonic decline of mutant numbers with the death-to-birth ratio requires the mutant production and the selective disadvantage to have equilibrated, which takes longer to occur for smaller selective disadvantages (*s*_2_).

To summarize, in an exponentially growing population, the effect of the death-to-birth ratio on deleterious mutant burden depends both on the parameter in which the disadvantage is expressed and on the threshold population size under consideration. The most relevant death-to-birth ratio ranges are those, for which the mutant cells can sustain their own growth. If the mutant is disadvantageous due to a reduced birth rate, deleterious mutant burden generally increases with the death-to-birth ratio in this parameter range, and this relationship tends to become less pronounced for larger threshold population sizes. If the mutant is disadvantageous due to an increased death rate, we can observe a decline of mutant burden with higher death-to-birth ratios, and this outcome becomes more pronounced for larger threshold population sizes. If the fitness disadvantage is expressed in both the division and death rate of the mutant, combinations of the above-described effects can be observed, as summarized in SI.1.

#### A stochastic, spatial colony

In the spatial stochastic model, individuals occupy sites on a two-dimensional *L* × *L* grid, with each site containing at most one individual. Wild types attempt divisions at rate *r*_w_ and mutants at rate *r*_m_ = *r*_w_(1 + *s*_1_), while death occurs at rates *d*_w_ and *d*_m_ = *d*_w_(1 − *s*_2_), respectively, leaving the site empty. Upon a division attempt, a site is chosen uniformly from the Moore neighborhood and the offspring is placed there if it is unoccupied; the division is aborted otherwise. Mutations arise only during wild-type division: with probability *µ* the offspring is mutant, otherwise it is wild type. These events occur probabilistically, see Methods Section “A stochastic, spatial system” for details. The system is initiated from a single wild type at the center of the grid and trajectories are followed until the total population size first reaches *N* ; the grid size is set large enough that the expanding population never reaches the boundary. Runs that end in complete population extinction before size *N* is reached are excluded. As before, we investigate how average mutant numbers depend on the death-to-birth parameter *δ*.

Similar to the non-spatial model described above, advantageous mutants always show an increase in numbers as the death-to-birth ratio increases. For disadvantageous mutants, we observe a fundamental change compared to the perfectly mixed, exponential growth scenario studied above. We again first explore the assumption that the mutant disadvantage is expressed by a reduced birth rate of cells, and then assume that the mutant disadvantage is expressed through a higher death rate of cells.

Figure 3(a) illustrates the case where the mutants are disadvantageous in birth. It shows the average number of mutants over many realizations of the agent-based model as a function of total population size (wild-type + mutant) in the spatially growing colony of cells. Different lines show such simulations for different values of the death-to-birth parameter, *δ*. At relatively low population sizes, we observe a similar trend as for the mass-action scenario: an increase in the death-to-birth ratio leads to a larger number of mutants. However, at high population sizes, this trend reverses, and now we observe a monotonic decline of mutant numbers as a function of the death-to-birth ratio. This is never observed for a mass-action system for the ranges of *δ* considered here. The hierarchy of the average number of mutants for different values of *δ* crosses over at intermediate population sizes, *N* . This is reflected in a one-humped relationship between mutant numbers and the death-to-birth ratio for those intermediate population sizes. The simulations in Figure 3(a) assume a relative large selective disadvantage of 10% (*s*_1_ = −0.1), and in this case, the crossover occurs for population sizes between 10^4^ and 10^5^. Importantly, however, the same general pattern is observed for smaller selective costs, although the cross-over occurs at higher population sizes in this case. While this becomes computationally more costly to simulate, we show in Figure SI.6(d) that for a 2% fitness cost (*s*_1_ = −0.02), the crossover happens at a total population size of around 10^6^. For even smaller selective disadvantages, the crossover will occur at larger populations sizes, which, however, will still be biologically relevant given that the population size of tumors may reach levels around 10^12^ cells or more.

**Figure 3:**
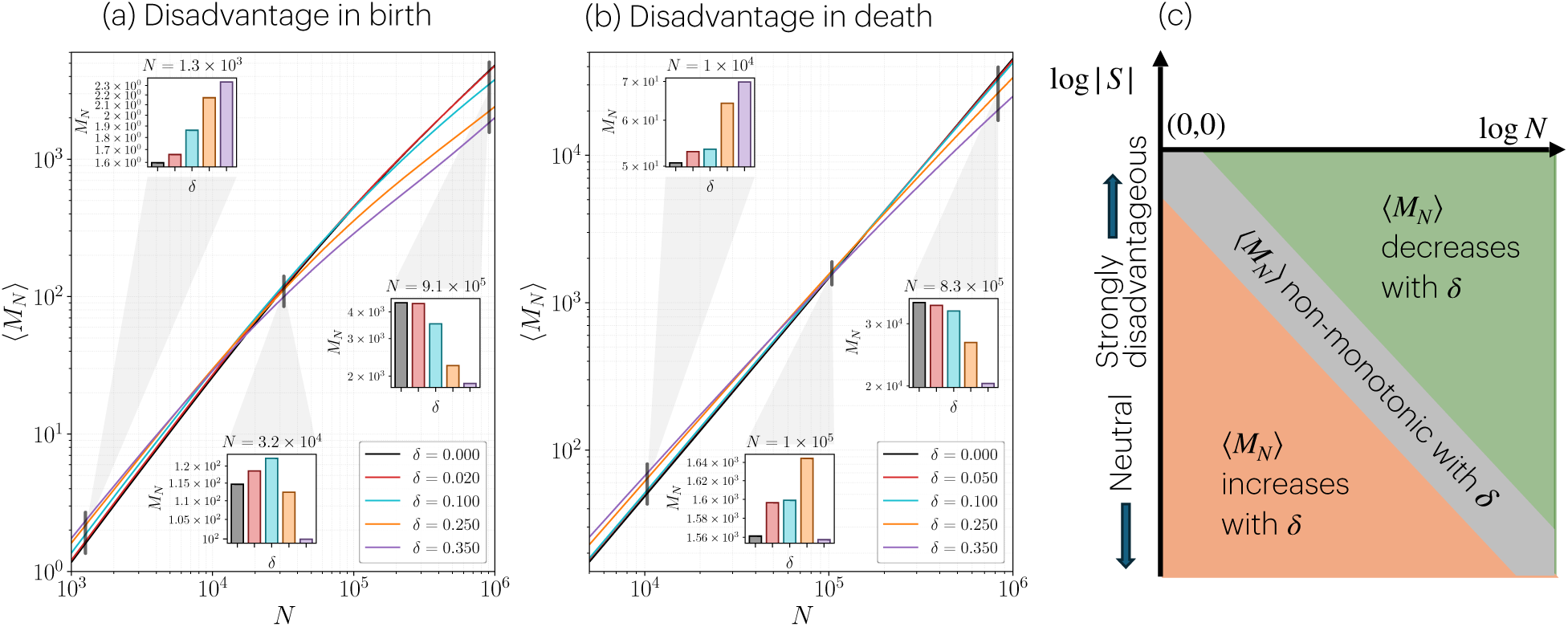
The spatial model: dependence of disadvantageous mutant burden on the population size for different death-to-birth ratios. (a) The expected number of mutants, ⟨*M_N_* ⟩, as a function of final population size *N* when disadvantage acts only through birth, (*s*_1_*, s*_2_) = (−0.10, 0.00). Different colors indicate different death-to-birth ratios *δ*, and the inset shows ⟨*M_N_* ⟩ across death-to-birth ratios at selected values of *N* . (b) Same as (a), but disadvantage acts only through death, (*s*_1_*, s*_2_) = (0.00, −0.05). In (a,b), each curve represents the average over 10^5^ independent simulations. Other parameters are *r*_w_ = 0.5 and *µ* = 10*^−^*^4^. (c) A schematic showing how the dependence of ⟨*M_N_* ⟩ on *δ* for disadvantageous mutants changes with |*s*| and *N*.

Although the relationship between mutant burden and the death-to-birth ratio appears complex at first glance, a simple and systematic pattern emerges, if we evoke spatial aspects of mutant and wild-type co-dynamics. Growth occurs within a narrow proliferative layer at the colony’s advancing edge. When a mutant arises near this front, its descendants form a wedge-like cluster whose boundaries shift through stochastic fluctuations [30–32]. Under low death-to-birth ratios and mild selective disadvantage, the mutant cluster can expand laterally, forming temporary sectors that elevate overall mutant burden above well-mixed expectations [29], see Figure 4(a). At higher death-to-birth ratios, frequent deaths not only diffuse the frontier and slow colony expansion, but also erode mutant clusters (Figure 4(b)), accounting for the negative correlation between mutant numbers and death-to-birth ratios at large population sizes. Note that the populations shown in Figure 4(a,b) have an identical cell count, although the high-death-to-birth ratio case appears spatially larger because increased death events generate more empty sites and reduce the average population density. To explain this in more detail, consider two competing timescales:

1. The time, *T_N_* (*N, δ, s*), for the population to reach the final size *N* ; this quantity grows both with *N* and *δ* and only depends weakly on *s*. A longer time until the colony reaches size *N* also correlates with a larger number of cell divisions during this time, and hence with a larger number of mutants produced.
2. The lifetime of a mutant cluster after it appears, *T*_cluster_(*N, δ, s*), which is set by selective disadvantage and the death-to-birth ratio. Stronger disadvantage makes clusters shrink faster in the wild-type bulk, and higher death-to-birth ratios create more birth–death events that further speed up their erosion. Thus, *T*_cluster_ decreases both with *δ* and |*s*|.

**Figure 4:**
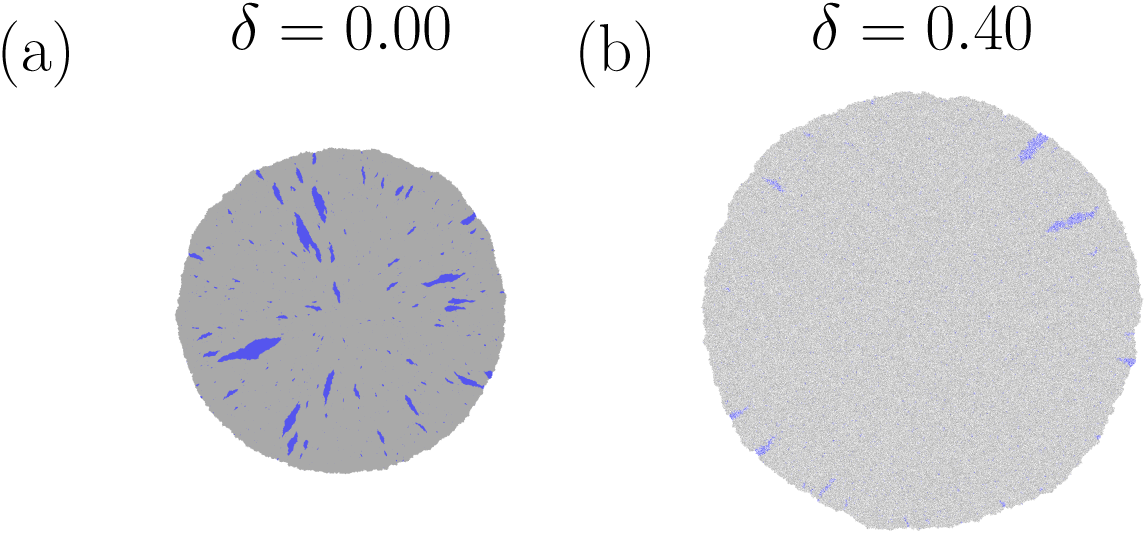
Dependence of the density of disadvantageous mutants bubble on increasing death-to-birth ratios. Final population snapshots for death-to-birth ratios (a) *δ* = 0 and (b) *δ* = 0.4, under *N* = 5 × 10^6^. Gray indicates the wild type and blue the mutant cells. Other parameters are *r*_w_ = 0.5, *µ* = 4 × 10*^−^*^4^, and (*s*_1_*, s*_2_) = (−0.05, 0.0). Note that the colony in (b) appears larger because of the decreased population density under *δ >* 0.

The death-to-birth ratio, *δ*, increases *T_N_* and decreases *T*_cluster_. For relatively small final population sizes, the time to reach *N* is shorter than the cluster lifetime (*T_N_ < T*_cluster_). Thus, higher death-to-birth ratios will generate more mutant sectors, while not enough time has passed for those disadvantageous mutant sectors to have been eroded by the wild-type. Hence, the number of mutants increases with the death-to-birth ratio. In the schematic of Figure 3(c), the left side of the phase diagram (smaller vales of *N*) corresponds to ⟨*M_N_* ⟩ increasing with *δ*.

As the threshold population size increases, so does *T_N_* . Once the growth timescale exceeds the sector survival timescale (*T_N_ > T*_cluster_), many sectors disappear before the population reaches size *N* . As a result, the typical cluster size decreases significantly with *δ*, which cannot be compensated by an increased number of clusters produced. This leads to an overall decrease in the mutant burden with *δ* at high *N* . In the schematic of Figure 3(c), the right side of the phase diagram (higher vales of *N*) corresponds to ⟨*M_N_* ⟩ decreasing with *δ*.

When the disadvantage is large, we never observe a crossover, and ⟨*M_N_* ⟩ decreases with *δ* (see Figure SI.6). As the selective disadvantage decreases, the survival time of mutant clusters increases, and consequently the crossover points shift to progressively larger final population sizes (the gray region in Figure 3(c) is moving to the right as we approach the neutral case). In the limit where the disadvantage tends to zero, the cluster survival timescale diverges, so the growth timescale never exceeds it within any finite population size; therefore, no crossovers are observed for neutral mutants (and *M_N_* always increases with the death-to-birth ratio at any *N*, see Figure SI.7).

The same mechanism applies if the mutant is assumed to be disadvantageous in death (see Figure 3(b)). In contrast to a disadvantage in birth, however, a decline of mutant numbers with the death-to-birth ratio at large threshold population sizes can already be seen in the well-mixed system, because the parameter *δ* multiplies the selection coefficient *s*_2_. The spatial dynamics described above are a conceptually different mechanism to result in a decline of mutant numbers at higher death-to-birth ratios, and here both effects act additively. Hence, the decline of mutant numbers with higher death-to-birth ratios is more pronounced if the disadvantage is expressed by a higher mutant death rate rather than a lower mutant birth rate.

To summarize (see the schematic in Figure 3(c)), the crossover (that is, a switch from a positive to negative *∂*⟨*N_M_* ⟩*/∂δ*) arises from competition between two timescales: the time *T_N_* required to reach size *N*, and the lifetime *T*_cluster_ of mutant clusters. Increasing the death-to-birth ratio raises mutation production but shortens cluster lifetime. When *T_N_ < T*_cluster_ (as is the case, for example, in figure 4(a)), clusters persist until size *N* is reached, so higher death-to-birth ratios increase mutant burden. When *T_N_ > T*_cluster_ (see figure 4(b)), clusters dissolve before the target size is reached, and higher death-to-birth ratios reduce mutant burden. The crossover occurs when *T_N_* ≈ *T*_cluster_. Because stronger selective disadvantage reduces cluster lifetime, the crossover shifts to smaller *N* as |*s*| increases.

#### Evolutionary rescue

So far, we have considered the average number of mutants as an evolutionary measure. Another outcome is the probability of evolutionary rescue, which is defined as the probability that a population survives an event that leads to the extinction of the wild-type, but leaves the mutants viable. In the context of growing populations, this is meaningful in parameter regimes in which the mutant burden is low enough to make survival of the population not guaranteed. Hence, we consider a scenario where the total cell population grows up to a threshold size, at which point all wild-type cells are eliminated and remaining mutants have a chance to grow. We determine the probability that the remaining mutants do not go extinct by stochastic effects, depending on the death-to-birth ratio of the cells (probability of rescue). We note that we focus on an extreme scenario where the adverse event immediately eliminates all wild-type cells, in order to study basic differences in spatially structured and well-mixed populations. A more comprehensive, future, study is required to investigate rescue dynamics in situations where the wild-type cell population decays at varying rates following the adverse event.

In the context of evolutionary rescue, we focus only on the case where mutants are disadvantageous before the onset of the adverse environment that subsequently wipes out the wild-type population. Specifically, advantageous mutants are not considered, since they can readily expand inside the bulk of the wild-type population before environmental deterioration, making evolutionary rescue almost inevitable. We observe fundamental differences in the dependence of the rescue probability on the death-to-birth ratio *δ* when comparing the well-mixed to the spatial system, assuming a mutant disadvantage in the birth rate (*s*_1_). In the well-mixed system, the rescue probability always declines with higher death-to-birth ratios (Figure. 5(a)), while in the spatial system, it can increase or show a non-monotonic dependence (Figure 5(d)).

**Figure 5:**
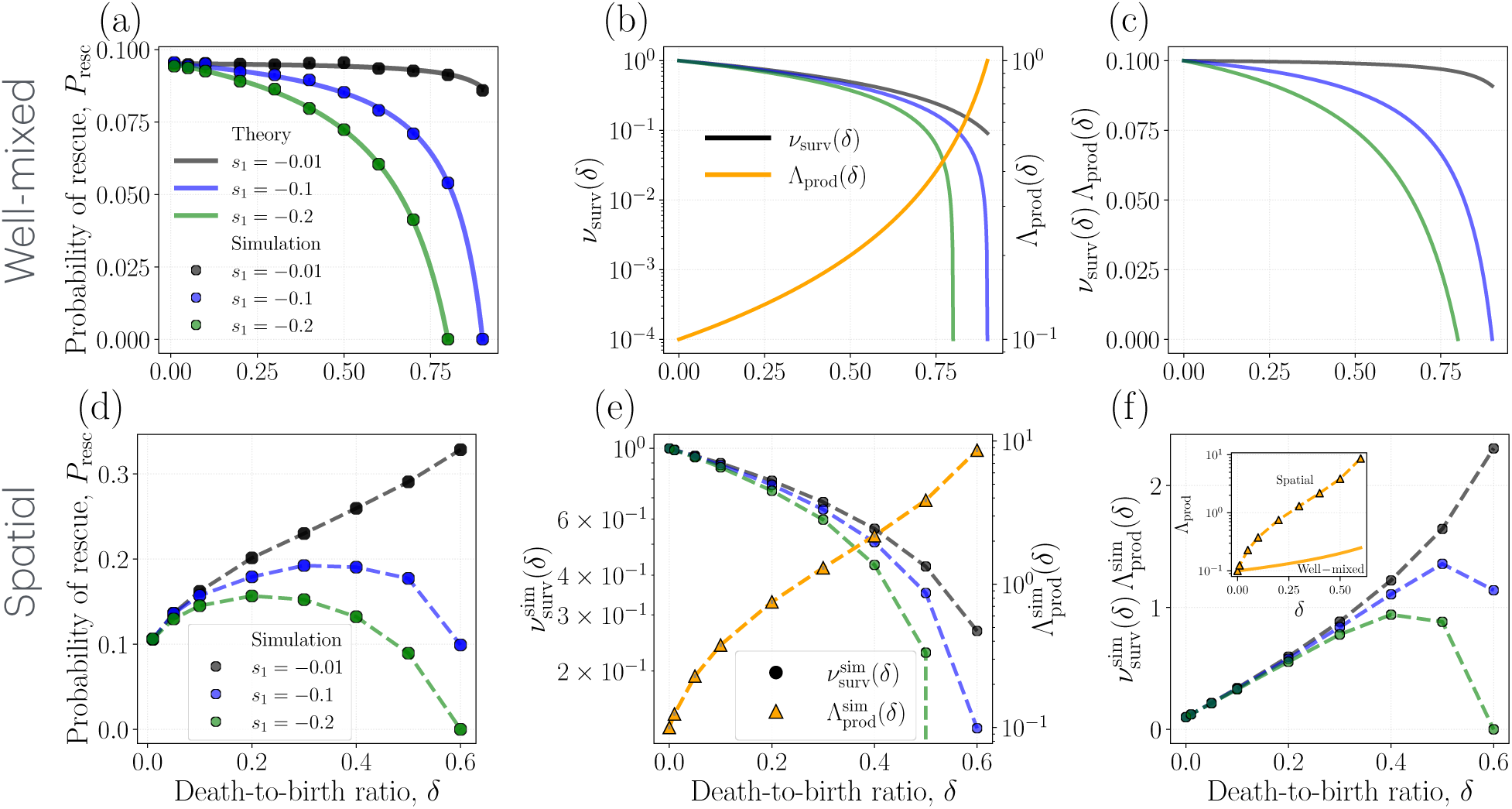
Rescue mechanism in well-mixed and spatial systems by disadvantageous mutants. We consider mutants that experience selective disadvantage only through birth, with no death disadvantage (*s*_2_ = 0). The mutation rate is fixed at *µ* = 10*^−^*^5^, the wild-type birth rate at *r*_w_ = 0.25, and all wild types are eliminated once the total population reaches *N* = 10^4^. Panels (a–c) show the well-mixed stochastic system, while panels (d–f) show the spatial system. Panels (a) and (d) display the rescue probability as a function of the death-to-birth ratio *δ* for different mutant disadvantages. Panels (b) and (e) show the single-lineage survival probability *ν*_surv_ together with the mutant production measure Λ_prod_. In the well-mixed system, increasing the death-to-birth ratio decreases *ν*_surv_ and increases Λ_prod_, but their product always decreases, leading to a monotonic decline in rescue probability. In contrast, the spatial system exhibits a much faster increase in mutant production with increasing death-to-birth ratios (see inset), which can compensate for the reduced survival probability and produce a non-monotonic, and even increasing, rescue probability for weakly disadvantageous mutants. Panels (c) and (f) show the product *ν*_surv_Λ_prod_, highlighting the distinction in rescue in the two systems. Simulation results are shown as symbols, with each data point averaged over at least 10^5^ independent realizations. Standard error bars are smaller than the symbol size.

In SI.2.2, we show analytically that in the well-mixed system, increasing the death-to-birth parameter *δ* always decreases the rescue probability, and that no other behavior is possible for disadvantageous mutants satisfying (*s*_1_ + *s*_2_) ≤ 0. Although the average number of mutants tends to increase for higher death-to-birth ratios (as we have shown above), this measure is not relevant for determining rescue probabilities, because average mutant numbers can be driven by jackpot type events that have no impact on rescue.

Instead, the theory in SI.2.2 points to two important quantities that govern the rescue dynamics. The first is the probability that a single mutant lineage survives and ultimately rescues the population, denoted by *ν*_surv_. The second is the total mutant production by the wild-type population before the onset of the adverse condition in which all wild types are eliminated, denoted by Λ_prod_. Importantly, Λ_prod_ is distinct from the mutant burden *M_N_* . While *M_N_* depends on the selective disadvantage of the mutant, Λ_prod_ is independent of mutant fitness and depends only on the mutation rate and the growth dynamics of the wild-type population. We theoretically compute the rescue probability (SI.2.2) in terms of the mutant survival probability *ν*_surv_ and the mutant production factor Λ_prod_ as

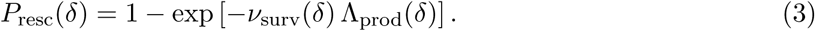

Here,

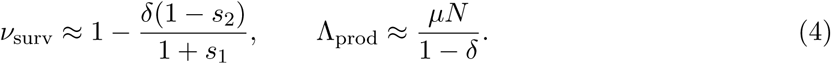

This theoretical estimate accurately captures the death-to-birth ratio dependence of the rescue probability, as demonstrated by the solid curves in Figure 5(a).

As the death-to-birth ratio increases, the survival probability decreases, and the decrease becomes faster for mutants with larger selective disadvantage. In contrast, mutant production increases with the death-to-birth ratio (see Figure 5(b)). The rescue probability is a monotonic function of the product of these two quantities, which, for disadvantageous mutants, always decreases with the death-to-birth ratio (see Figure 5(c)).

This pattern is different for the spatial system (Figure 5(d)). For relatively small mutant disadvantage (*s* = −0.01), the rescue probability increases monotonically with the death-to-birth ratio *δ* (see the black symbols in Figure 5(d)). We computed the survival probability 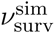 from simulations by introducing a single mutant and measuring the probability that its lineage reaches the target population size. Similar to the well-mixed system, this survival probability decreases with increasing death-to-birth ratios, and the decrease becomes sharper for mutants with larger selective disadvantage (Figure 5(e)). We also find that the mutant production generated by the wild-type population, measured from simulations as 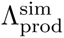, increases with death-to-birth ratios (see the yellow symbols in Figure 5(e)).

Interestingly, in the spatial model, mutant production increases much faster with death-to-birth ratios than in the well-mixed model (see the inset of Figure 5(f)). This creates a competition between two opposing effects. On one hand, increasing death-to-birth ratios reduce the probability that an individual mutant lineage survives and rescues the population. On the other hand, higher death-to-birth ratios simultaneously generate mutants at a much higher rate, substantially increasing the number of independent mutant lineages produced before the adverse condition begins. For weakly disadvantageous mutants, the enhanced mutant production can compensate for the reduced single-lineage survival probability and unlike the well-mixed case, where the product decreases monotonically with *δ*, the spatial system with weakly disadvantageous mutant can exhibit an increase in the product of the survival probability and mutant production (see the black symbols in Figure 5(f)). Consequently, the rescue probability can increase with the death-to-birth ratio within the observed range of values.

As the strength of disadvantage increases, mutant production can still compensate for the reduction in lineage survival probability at small death-to-birth ratio values, leading to an initial increase in rescue probability. However, at larger death-to-birth ratios, mutant production is no longer sufficient to offset the decline in survival probability, causing the rescue probability to decrease with further increases in the death-to-birth ratio (see the blue and green symbols in Figure 5(d)).

Although directly substituting the product of the lineage survival probability and mutant production estimated for the spatial system into the well-mixed rescue formula does not yield an exact prediction for the spatial system, the failure of mutant production to compensate for the reduced lineage survival probability at large death-to-birth ratios is clearly reflected in the behavior of their product for intermediate and strongly disadvantageous mutants (see the blue and green symbols in Figure 5(f)).

We note that, for sufficiently large death-to-birth ratios, the survival probability of mutant lineages becomes vanishingly small. Consequently, despite the enhanced mutant production, the rescue probability must eventually decrease toward zero for all mutants, including weakly disadvantageous ones. Due to the computational infeasibility of simulating very large death-to-birth ratios, we are unable to directly observe this eventual decrease in rescue probability for weakly disadvantageous mutants.

## 3 Discussion

While patterns of mutant burden in expanding colonies have been analyzed with stochastic birth death processes, the dependence of mutant burden on cellular death-to-birth ratios has not been explicitly and systematically investigated so far, both in well-mixed and spatially structured populations. For well-mixed populations, the argument has been made that higher death-to-birth ratios lead to a higher mutant burden, because more cell divisions are required to reach the threshold population size, which correlates with more mutant generation [16]. While this particular dependence can be correct (and is the only dependence for advantageous mutants), a systematic exploration for disadvantageous mutants has so far been missing. Yet, an understanding of how deleterious mutant burden correlates with the cellular death-to-birth ratio can be biologically important for interpreting the potential for populations to evolve, and tumor evolution is a prominent example of this. Tumors progress via multi-stage evolutionary processes [33, 34], where mutant clones that emerge allow tumor cells to adapt to their microenvironment, leading to a more efficient growth, the ability to overcome selective barriers, and hence to accelerated disease progression. Sometimes, mutants that are deleterious in the current environment can gain an advantage following an environmental change, with drug resistance being a well-known example of this, see e.g. [35, 36] in the context of bacterial drug resistance and [37–39] in the context of cancer drug resistance. Furthermore, a higher number of clones with a single mutational hit increases the chances that one of these single-hit mutants can accumulate further mutations. This enables the evolution of more complex cell phenotypes, where a multi-hit mutant has a selective advantage while the intermediate mutants are deleterious [40–42]. Such crossing of fitness valleys is thought to be a process that contributes to tumor progression e.g. in the context of tumor suppressor gene inactivation [43–45]

Tumor cell death-to-birth ratios not only vary across different cancers, but they can also vary within the same type of cancer across different patients, and can further change over time as cancer progresses. Clinical studies have investigated the relationship between tumor cell death-to-birth ratios and prognosis / survival outcomes. While some studies have shown that higher death-to-birth ratios correlate with poor outcomes [46–49], other studies have demonstrated the opposite [26, 27], resulting in contradictory observations. Our analysis, however, has shown that either trend is possible, depending on the parameter regions and details of the population structure. We have shown that both low and high death-to-birth ratios, or even intermediate death-to-birth ratios can maximize the deleterious mutant burden, and hence the degree of standing genetic variation that can promote the ability of the tumor to adapt to micro-environmental change. These results thus enable a more informed interpretation of clinical data.

Our analysis has highlighted two factors that have a major influence on the relationship between mutant burden and the death-to-birth ratio. (i) First, population structure plays a crucial role. If the mutant fitness cost is expressed as a reduced division rate, then a higher death-to-birth ratio always leads to a larger mutant burden for the well-mixed system in the range of *δ* values for which the mutant is viable. With spatial population structure, in contrast, this relationship reverses as the population grows, such that higher cell death-to-birth ratios lead to a lower burden of disadvantageous mutants. Our analysis indicates that this can occur for relatively small selective disadvantages (e.g. 1%) at realistic population sizes that are well below those of detectable tumors. (ii) The second factor is the demographic parameter in which the mutant fitness cost is expressed. If it is expressed through an increased mutant death rate compared to the wild-type, then the fitness disadvantage, *s*_2_, multiplies the death-to-birth parameter *δ*, resulting in an increased impact of the fitness cost for higher death-to-birth ratios. This can lead to a negative correlation between mutant burden and the death-to-birth ratio for well-mixed populations, especially if the selective disadvantage is relatively large, and this effect is not observed if the mutant is disadvantageous through a reduced division rate. We note that this effect is separate from the spatial dynamics (which can independently lead to a negative correlation between mutant burden and the death-to-birth ratio). In contrast to the spatial dynamics, however, the fitness cost of the mutant needs to be relatively large to observe this effect.

In the spatial dynamics, we note that in the limit of large populations, a higher death-to-birth ratio always leads to a lower burden of deleterious mutants. However, at lower population sizes, the opposite can be observed and higher death-to-birth ratios lead to more deleterious mutants. For intermediate population sizes, we can observe a non-monotonic relationship. The lower the fitness cost of a mutant, the larger is the population size at which the relationship between the death-to-birth ratio and mutant burden reverses to being negative. This is another insight that can be useful when interpreting biological data.

We also studied the effect of the cell death-to-birth ratio on evolutionary rescue, i.e. the probability of mutants to ensure survival of the cell colony following an environmental change that drives the wild-type cell population extinct. This becomes relevant if the number of mutants in the cell population is sufficiently low, such that survival of the existing mutants is not guaranteed. This would be less relevant to established tumors, where the population size is large relative to the inverse mutation rate, but is applicable to smaller populations that experience a catastrophic environmental change. In the current study, we concentrated on the scenario where the catastrophic event instantly eliminates the wild-type population, to study basic differences in rescue probabilities in well-mixed and spatially structured populations. Here, the average number of mutants at a given population size is less important, but the probability to have no mutants or a few mutants is an important determinant of rescue. We showed that the rescue probability displays different properties compared to the mutant burden at a threshold population size, and again that spatial structure can substantially change outcome. For well-mixed systems, the rescue probability always declines with the death-to-birth ratio, because the reduced ability of mutant clones to grow and establish themselves is the dominant contributor here. For spatially structured populations, however, it is possible that the rescue probability increases with the death-to-birth ratio, or that we observe a non-monotonic dependence. The reason is that with spatial structure, increased mutant production at higher death-to-birth ratios plays a more dominant role compared to the reduced cell clone survival probability. Hence, population structure again is a crucial factor that determines how evolutionary outcome is influenced by the death-to-birth ratio, with important implications for the interpretation of data. In future work, it will be important to study how these patterns change if the wild-type cell population is not immediately extinguished when the environment changes. The prolonged persistence of wild-type cells on their way to extinction can lead to further mutant generation events, which can contribute to rescue. Additionally, for spatially structured populations, the prolonged presence of the wild-type population can inhibit the mutant cell population to grow, similar in principle to arguments made in the context of adaptive therapy of cancer [50, 51].

Model predictions presented here could be tested using microbial fluctuation assays and spatial colony expansion experiments that quantify mutant clone distributions in well-mixed and spatial populations (similar to the experiments in [52–54]). Further, experimental evolution studies could be done with rescue systems in which populations are exposed to severe environmental stress and rescue frequencies are measured across replicate populations [55]. Although the present work focuses on two-dimensional systems, we expect the qualitative behavior to remain robust in three dimensions. Quantitative differences may arise due to faster effective growth and the larger number of neighboring cells contributing to local dynamics. Investigating these effects explicitly in three-dimensional systems represents an important direction for future work.

We discussed many of our theoretical results in the context of carcinogenesis and cancer evolution, because the death-to-birth ratio clearly varies among different cancers, among different individuals who have the same cancer, and even within individuals as the tumor evolves. Having a comprehensive understanding of how the cell death-to-birth ratio affects evolutionary processes in expanding populations allows a better interpretation of clinical data that otherwise would seem contradictory. The theoretical insights developed here, however, have more general implications for evolutionary processes in asexual populations, whether this is bacteria or other lower organisms that need to adapt to changing environments. The presence of currently deleterious mutants often helps a population to survive following an environmental change, or serves as an evolutionary substrate for the evolution of more complex phenotypes. The work presented here helps us understand more completely how the death-to-birth ratio, which is a basic demographic characteristic of reproducing organisms, influences the ability of populations to adapt, evolve, and survive.

## 4 Methods

### 4.1 A stochastic, well-mixed system

Within the well-mixed stochastic model, let *X*(*t*) denote the number of wild types at time *t*, and *Y* (*t*) the number of mutants. We use capital letters, in contrast to the deterministic well-mixed system, to emphasize the stochasticity and discreteness of these variables. The stochastic system evolves via five distinct types of reactions. A wild-type individual reproduces at a rate proportional to *r*_w_. Each reproduction event yields either a wild-type offspring, with probability 1 − *µ*, or a mutant offspring, with probability *µ*. Wild-type individuals die at a rate proportional to *d*_w_. Mutants reproduce at a rate proportional to *r*_m_ and die at a rate proportional to *d*_m_. The dynamics can be summarized by the following five reactions:

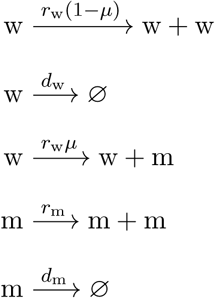

Equivalently, the dynamics of the stochastic variables *X*(*t*) and *Y* (*t*) can be written as

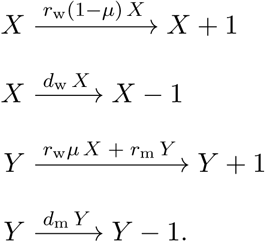

We simulate the stochastic dynamics starting from the initial condition (*X*(0)*, Y* (0)) = (1, 0). The system admits an absorbing state at (0, 0), which we exclude from our analysis. Thus, all results for the stochastic well-mixed system are obtained from trajectories that terminate upon first hitting the target size *X* + *Y* = *N* . Specifically, for each independent realization, the number of mutants *M_N_* at the hitting time is recorded, and the ensemble-averaged mutant burden ⟨*M* ^stoch^⟩ is computed.

### 4.2 A stochastic, spatial system

To simulate the stochastic dynamics of the spatial colony, we implemented a discrete event-based algorithm on a two-dimensional lattice, where each lattice site can either be empty, occupied by a wild-type cell, or occupied by a mutant cell. The lattice has size *L* × *L*, large enough to ensure that the boundary of the expanding population never reaches the boundary of the lattice, and initially only the central site is occupied by a single wild-type cell. At each update, a randomly chosen occupied site (*i, j*) is selected from the list of all occupied positions. If the selected cell is wild type, it dies with probability *d*_w_ or attempts to divide with probability *r*_w_; if it divides, one of the eight neighboring sites (Moore neighborhood) is chosen uniformly at random. If the target site is empty, the offspring is placed there; otherwise, the division is aborted. Mutation occurs only during the division of a wild-type cell: With probability *µ* the offspring becomes mutant, and with probability 1 − *µ* it remains wild type, corresponding to the reactions

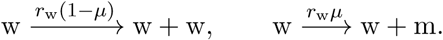

If the selected cell is mutant, it dies with probability *d*_m_ = *d*_w_(1 − *s*_2_) or divides with probability *r*_m_ = *r*_w_(1 + *s*_1_), producing a mutant offspring when an empty neighboring site is available and chosen uniformly and randomly to place the offspring, according to

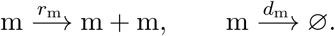

Each event updates the grid and the population counts (*X, Y*), where *X* and *Y* denote the numbers of wild-type and mutant cells, respectively. The simulation proceeds asynchronously, with on average one update attempt per occupied site per iteration, until a stopping condition is reached. The process is terminated either when the total population size first reaches the predefined target *X* + *Y* = *N* or when the population goes extinct (*X, Y*) = (0, 0). In the latter case, the system is reinitialized with a single wild-type cell at the center, and the run is repeated. For each independent realization, the number of mutants *M_N_* at the hitting time *X* + *Y* = *N* is recorded, and the ensemble-averaged mutant burden ⟨*M* ^spatial^⟩ is obtained from all non-extinct trajectories.

## Acknowledgments

SSM gratefully acknowledges fruitful discussions with Ramanarayanan Kizhuttil and Sydney Acker-mann.

## Author contributions

N.L.K., and D.W. designed research; S.S.M., N.L.K., and D.W. performed research; S.S.M., N.L.K., and D.W. analyzed the results; and S.S.M., N.L.K., and D.W. wrote the paper.

## Data and code availability

The authors affirm that all data necessary to confirm the conclusions of this article are presented within the main text and figures. Supplemental information and representative codes are available at https://github.com/samratsohel/mutant-burden.git.

