## Supplemental Materials for "Spatial population structure can reverse the dependence of mutant burden on the death-to-birth ratio of cells"

### Supplementary Information

#### SI.1 A deterministic, well-mixed colony.

In order to provide some simple intuition for the stochastic systems studied here, we will list some properties of the corresponding non-spatial, deterministic system given by the following ordinary differential equations:

$$\dot{x} = r_{\mathrm{w}} x(1-\mu) - d_{\mathrm{w}} x, \tag{1a}$$

$$\dot{y} = \mu r_{\mathrm{w}} x + r_{\mathrm{m}} y - d_{\mathrm{m}} y. \tag{1b}$$

where $x(t)$ and $y(t)$ measure the populations of wild-type and mutant cells respectively. Under the initial condition $x(0) > 0,\ y(0) = 0$, we let the population grow to the total

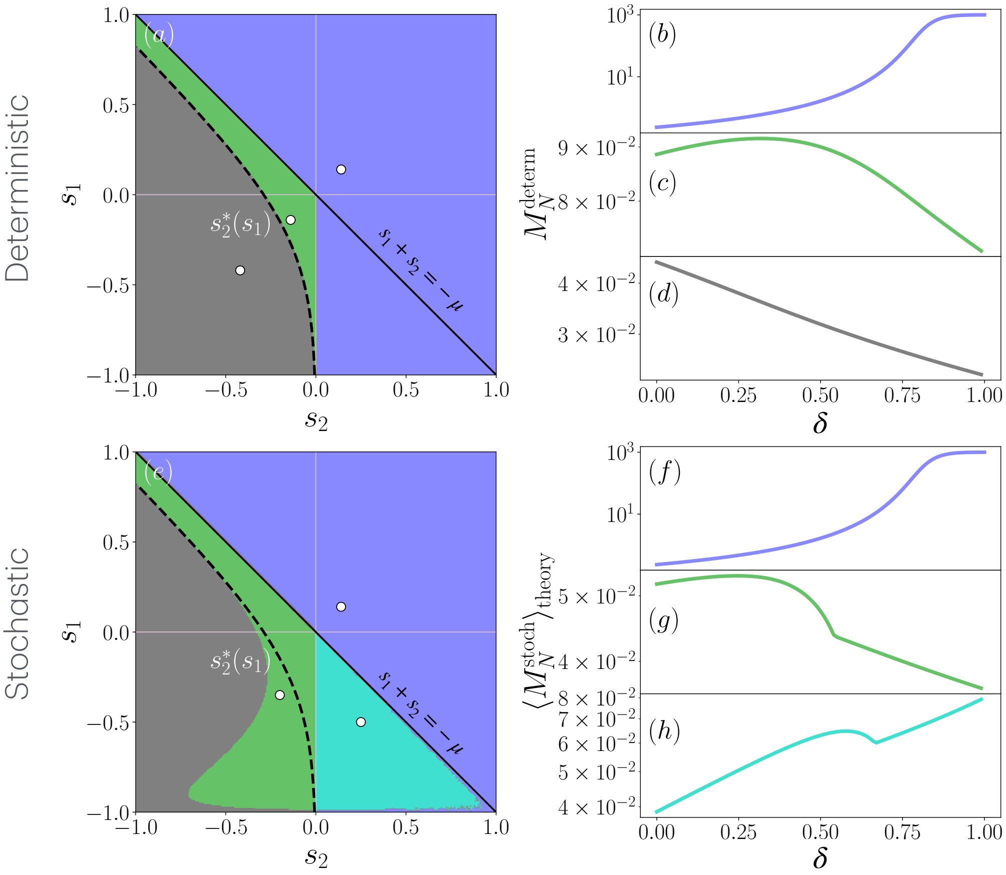

Figure SI.1: *Dependence of mutant burden on the death-to-birth ratio in well mixed systems.* (a)–(d) represent outcomes from the deterministic model, and (e)–(h) those from the stochastic model (using Equation (20)). (a) and (e) are phase diagrams showing the qualitative behavior of mutant accumulation across the $(s_1, s_2)$ parameter space for the deterministic and stochastic models, respectively: Blue indicates a monotonic increase; green denotes non-monotonic behavior with a decrease in mutant accumulation at high death-to-birth ratios; gray indicates a monotonic decrease; and teal represents non-monotonic behavior with an increase at large death-to-birth ratios. The line $s_2^*(s_1) = (s_1^2 \ln N)/(1 - N^{-s_1} - s_1 \ln N)$ (see SI.1.4 for a derivation) separates the green and gray regions in the deterministic case. (b)–(d) show illustrative mutant accumulation curves as functions of the death-to-birth ratio $\delta$, corresponding to the white dots marked in (a), which lie in the blue, green, and gray regions, respectively. These points correspond to $(s_1, s_2) = (0.14, 0.14)$, $(-0.14, -0.14)$, and $(-0.42, -0.42)$. Similarly, (f)–(h) display mutant accumulation curves as functions of $\delta$, corresponding to the white dots marked in (e), located in the blue, green, and teal regions, respectively, at $(s_1, s_2) = (0.14, 0.14)$, $(-0.35, -0.20)$, and $(-0.50, 0.25)$. The rest of the parameters are $r_{\mathrm{w}} = 1$, $\mu = 2 \times 10^{-5}$, and $N = 10^3$.

size $N$, and evaluate the mutant burden, $M_N$, as a function of the death-to-birth ratio, $\delta$. Figure SI.1(a-d) provides a summary of dependency of mutant burden on the death-to-birth ratio.

First, we note that advantageous mutant burden always increases with the death-to-birth ratio (see SI.1.2). Next, suppose first that mutants are deleterious because their division rate is smaller than that of the wild types ($s_1 < 0$), while their death rate is the same as that of the wild-type cells ($s_2 = 0$). In this case, again the mutant burden $M_N$ always increases with the death-to-birth ratio (see SI.1.3 for a detailed proof). This intuitive result is usually explained by noting that in a higher death-to-birth ratio colony, more cell divisions are required for the population to reach the target size $N$, and therefore there are more opportunities for mutant lineages to arise. This reasoning is not entirely correct and, in fact, breaks down when mutants are disadvantageous in terms of death.

Figure SI.1 shows three possible types of dependence on the death-to-birth ratio $\delta$: The number of mutants can increase with $\delta$, decrease, or vary in a non-monotonic way. For example, in the special case $s_1 = 0$ and $s_2 < 0$, where mutants are disadvantaged only in terms of death, the mutant burden as a function of the death-to-birth ratio, $M_N(\delta)$, first increases and then decreases with $\delta$ for small values of $|s_2|$. Moreover, as $|s_2|$ exceeds a threshold $|s_2^*|$ (see SI.1.4 for a detailed proof), the function becomes monotonically decreasing in $\delta$. A significant decay of $M_N$ with $\delta$ is observed when the selection coefficient $|s_2|$ exceeds $\sim 1/\ln N$ (see SI.1.5 for a detailed proof).

In Figure SI.1(a,e), there are two lines that are important in terms of mutant vs wild-type net growth rates. Consider the net wild-type growth rate, $G_{\mathrm{w}} = r_{\mathrm{w}}(1-\mu) - d_{\mathrm{w}}$, and the mutant net growth rate, $G_{\mathrm{m}} = r_{\mathrm{m}} - d_{\mathrm{m}}$. The two are equal when

$$D(\delta) \equiv G_{\mathrm{w}} - G_{\mathrm{m}} = -r_{\mathrm{w}}(s_1 + \delta s_2 + \mu), \tag{2}$$

is zero:

$$(s_1 + \delta s_2 + \mu) = 0.$$

The difference between the two growth rates is $\delta$-dependent, and the two extreme cases correspond to $\delta = 0$ and $\delta = 1$. These two cases give rise to the lines $s_1 = \mu$ and $s_1 + s_2 = -\mu$ (the former line is very close to the $s_1 = 0$ in Figure SI.1(a,e) and is not marked separately). In particular, the set of $(s_1, s_2)$ values that is below both of these lines corresponds to the parameter sets where the mutant is disadvantageous (grow slower) for all values of $\delta$. In this region we observe only green and gray shading in both panels (a) and (e) of figure SI.1 (that is, the mutant burden decreases with $\delta$ as $\delta$ approaches 1). Outside this region, the shading is blue or teal, that is, the mutant burden increases with $\delta$ as $\delta$ approaches 1. More details are found in the next sections.

#### SI.1.1 Reformulation of deterministic well-mixed model

The parameter $\mu$ is a small non-negative constant, and the condition $\delta \in [0, 1-\mu)$ guarantees exponential growth of the wild-type subpopulation; that is, in this model, the wild type is always supercritical. The selective advantage parameters $s_1$ and $s_2$ are assumed to lie within the interval $[-1, 1]$. The system can be reformulated in a more compact form by rescaling time as $t^* = r_{\mathrm{w}} t$, which yields

$$\frac{dx}{dt^*} = (1 - \delta - \mu)\, x, \tag{3a}$$

$$\frac{dy}{dt^*} = \mu x + \big[(1-\delta) + (s_1 + \delta s_2)\big] y. \tag{3b}$$

With the total population size $T = x + y$ and the mutant frequency $p = y/N$, the system can be rewritten as

$$\frac{dN}{dt^*} = \bar{r}(t^*)\, N, \tag{4a}$$
$$\frac{dp}{dt^*} = s\, p(1-p) + \mu(1-p), \tag{4b}$$

where the effective selection coefficient and mean Malthusian fitness are given by

$$s = s_1 + \delta s_2, \tag{5}$$
$$\bar{r}(t^*) = (1 - \delta - \mu) + s\, p(t^*). \tag{6}$$

Starting from the initial condition $p(0) = 0$, it admits the explicit solution

$$p(t^*) = \frac{1 - e^{-(s+\mu)t^*}}{1 + \frac{s}{\mu} e^{-(s+\mu)t^*}}. \tag{7}$$

where $\alpha = s + \mu$ sets the characteristic rate of approach to equilibrium. The total population size evolves according to the time-dependent Malthusian growth rate $\bar{r}(t^*)$, and the time required to reach a prescribed population size $N$ is defined implicitly by the condition

$$N(t_N^*) = N, \tag{8}$$

The number of mutants at this stopping time is therefore given by

$$M_N = N\, p(t_N^*). \tag{9}$$

##### SI.1.2 Advantageous mutant burden increases with the death-to-birth ratio

We consider advantageous mutants, defined by $s_1 > 0$ and $s_2 > 0$, and moreover $\partial_\delta s = s_2 > 0$. For any fixed $\delta$ the mutant frequency $p(t^*)$ evolves as

$$\frac{dp}{dt^*} = (1-p)(sp + \mu). \tag{10}$$

Since $0 \leq p \leq 1$, $s > 0$, and $\mu > 0$, it follows that $(1-p) > 0$ and $(sp + \mu) > 0$, hence

$$\frac{dp}{dt^*} > 0 \quad \forall t^* > 0. \tag{11}$$

Thus, the mutant frequency increases monotonically with time. At fixed time $t^*$, the frequency depends on $\delta$ through $s = s_1 + \delta s_2$. Since $s_2 > 0$, increasing the death-to-birth ratio increases $s$. From the explicit solution

$$p(t^*) = \frac{1 - e^{-(s+\mu)t^*}}{1 + \frac{s}{\mu} e^{-(s+\mu)t^*}}, \tag{12}$$

it follows that $p(t^*)$ is an increasing function of $s$, and therefore

$$\frac{\partial p}{\partial \delta} > 0 \quad \forall t^* > 0. \tag{13}$$

We now compare two close death-to-birth ratio values, $\delta$ and $\delta + d\delta$. The time required to reach a fixed population size $N$ is defined implicitly by

$$N(t_N^*) = N. \tag{14}$$

Since increasing the death-to-birth ratio reduces the net growth rate, the time to reach size $N$ increases, so that

$$t_N^*(\delta) < t_N^*(\delta + d\delta). \tag{15}$$

At the earlier time $t_N^*(\delta)$, the system with larger death-to-birth ratio already has a higher mutant frequency because $\partial_\delta p > 0$. Furthermore, the additional time required to reach the same population size further increases the mutant frequency because $\partial_{t^*} p > 0$. Combining these two effects, we conclude that for advantageous mutants,

$$M_N = N\, p(t_N^*) \tag{16}$$

is an increasing function of the death-to-birth ratio $\delta$.

##### SI.1.3 Condition for the decrease of $M_N$ with the death-to-birth ratio at $\delta \to 1^-$

To obtain a decreasing trend in mutant burden with the death-to-birth ratio near $\delta \to 1^-$, the mutant must be disadvantageous. This corresponds to a negative $D(\delta)$ (equation (2)), and in particular at $\delta \to 1^-$, we obtain

$$s_1 + s_2 < -\mu. \tag{17}$$

In this case, the mutant frequency remains small, $p(t^*) \ll 1$, so that the total population is dominated by the wild type. Hence, the population growth is approximately governed by the wild-type growth rate,

$$\frac{dN}{dt^*} \approx (1 - \delta - \mu)\, N. \tag{18}$$

Solving this equation gives

$$N(t^*) = x_0 \exp\big[(1 - \delta - \mu) t^*\big]. \tag{19}$$

The time required to reach population size $N$ is therefore

$$t_N^* \approx \frac{1}{1-\delta} \log\left(\frac{N}{x_0}\right). \tag{20}$$

For small $p$, the frequency dynamics reduce to

$$\frac{dp}{dt^*} \approx s\, p + \mu. \tag{21}$$

Solving above equation with initial condition $p(0) = 0$ gives

$$p(t^*) = \frac{\mu}{|s|}\left(1 - e^{-|s|t^*}\right). \tag{22}$$

As $\delta \to 1^-$, we have $1 - \delta \to 0$, and therefore $t_N^* \to \infty$. In this limit,

$$p(t_N^*) \to \frac{\mu}{|s|}. \tag{23}$$

Hence, the mutant number at size $N$ is

$$M_N = N\, p(t_N^*) \approx N \frac{\mu}{|s|} = \frac{N\mu}{-(s_1 + \delta s_2)}. \tag{24}$$

Differentiating with respect to $\delta$ gives

$$\frac{dM_N}{d\delta} = \frac{N\mu\, s_2}{(s_1 + \delta s_2)^2}. \tag{25}$$

Evaluating near $\delta \to 1^-$,

$$\left.\frac{dM_N}{d\delta}\right|_{\delta \to 1^-} = \frac{N\mu\, s_2}{(s_1 + s_2)^2}. \tag{26}$$

Since the denominator is positive, the sign of the derivative is determined by $s_2$. Therefore,

$$\frac{dM_N}{d\delta} < 0 \quad \Longleftrightarrow \quad s_2 < 0. \tag{27}$$

Thus, a decrease in mutant burden with increasing death-to-birth ratio near $\delta \to 1^-$ requires that the disadvantage acts through increased death, i.e., $s_2 < 0$. If $s_2 \geq 0$, the mutant burden cannot decrease with the death-to-birth ratio in this limit.

##### SI.1.4 Analytic separator $s_2^\star(s_1)$ between green and gray regions in SI.1(a)

In terms of the original (unscaled) time variable, the net growth-rate difference between mutant and wild type is given by $D(\delta)$, see equation (2). Using the definition of the mutant number at the hitting time, we write

$$M_N(\delta) = N\, p(t_N), \tag{28}$$

where $p(t)$ denotes the mutant frequency. In the case where the mutant frequency remains small, using the mutation–selection dynamics we obtain the approximation

$$p(t) \approx \frac{\mu}{|s + \mu|}\left(1 - e^{-|s+\mu|t}\right). \tag{29}$$

Substituting this expression into the definition of $M_N(\delta)$, and using the relation between $s$ and $D(\delta)$, we obtain

$$M_N(\delta) \approx N \frac{r_{\mathrm{w}}\mu}{D(\delta)}\left[1 - \exp\big(-D(\delta)\, t_N(\delta)\big)\right]. \tag{30}$$

Here we present a derivation of the curve $s_2^\star(s_1)$ that separates parameter pairs $(s_1, s_2)$ into two distinct behaviors of the mutant accumulation $M_N(\delta)$: (i) a *non-monotonic* accumulation (the green region in FIG. SI.1(a), where $M_N(\delta)$ first increases for small $\delta$ and then decreases as $\delta \to 1^-$), and (ii) a *monotone-decreasing* accumulation (the gray region in FIG. SI.1(a)). The key observation is that $M_N(\delta)$ is the product of two factors: The steady-state prefactor $\frac{N r_\mathrm{w} \mu}{D(\delta)}$, which decreases with $\delta$, and the transient term $\left[1-\exp(-D(\delta)\, t_N(\delta))\right]$, which increases with $\delta$. Because one factor decreases while the other increases, the product can have at most a single interior extremum (a maximum). For such a maximum to occur, the initial relative growth of the transient term must outweigh the initial relative decay of the steady-state prefactor, equivalently, $t_N \ll |D|^{-1}$. If this balance is not satisfied already at $\delta = 0$ (in particular, if $t_N(0) \gg |D(0)|^{-1}$), then the transient contribution never grows rapidly enough to compensate and $M_N(\delta)$ decreases monotonically. Therefore, the analytic separator between the green and gray regions is obtained by imposing the zero-slope condition at $\delta = 0$,

$$\left.\frac{\partial M_N}{\partial \delta}\right|_{\delta=0} = 0, \tag{31}$$

which yields the functional relation $s_2 = s_2^\star(s_1)$ distinguishing the two regimes. The object of interest is the composite function $M_N(\delta)$ obtained by substituting the $\delta$-dependent net disadvantage and the target-size hitting time into the closed-form expression for the mutant count at the time when the total population first reaches $N$,

$$M_N(\delta) = N \frac{r_\mathrm{w}\mu}{D(\delta)} \left[1 - \exp\big( - D(\delta)\, t_N(\delta)\big)\right]. \tag{32}$$

Differentiating $M_N(\delta)$ with respect to $\delta$ gives

$$\frac{\partial M_N}{\partial \delta} = N r_\mathrm{w} \mu \frac{\partial}{\partial \delta} \left( \frac{1}{D} \left[1 - e^{-D t_N}\right] \right). \tag{33}$$

Using the product and chain rules,

$$\begin{aligned} \frac{\partial M_N}{\partial \delta} = N r_\mathrm{w} \mu \Bigg[ & -\frac{D'}{D^2} \left(1 - e^{-D t_N}\right) \\ & + \frac{e^{-D t_N}}{D} \left(D' t_N + D t_N'\right) \Bigg], \end{aligned} \tag{34}$$

where derivatives with respect to $\delta$ are

$$\begin{aligned} D' &= \frac{\partial D}{\partial \delta} = -r_\mathrm{w} s_2, \\ t' &= \frac{\partial t_N}{\partial \delta} = \frac{\log(N/x_0)}{r_\mathrm{w}(1-\delta-\mu)^2}. \end{aligned} \tag{35}$$

Introduce for convenience the positive constant $C := \log(N/x_0)$. Next, we evaluate the relevant quantities at $\delta = 0$ as

$$\begin{aligned} D(0) &= -r_\mathrm{w}(s_1 + \mu), \\ t_N(0) &= \frac{C}{r_\mathrm{w}(1-\mu)}, \\ t_N'(0) &= \frac{C}{r_\mathrm{w}(1-\mu)^2}. \end{aligned} \tag{36}$$

The exponential factor $E := e^{-Dt_N}$ at $\delta = 0$ is

$$\begin{aligned} E(0) &= \exp\big(-D(0)t_N(0)\big) \\ &= \exp\left[\frac{C}{1-\mu}\,(s_1+\mu)\right]. \end{aligned} \tag{37}$$

So we can rewrite the aforementioned condition as

$$\left.\frac{\partial M_N}{\partial \delta}\right|_{\delta=0} = N r_{\rm w}\mu\left[-\frac{r_{\rm w}s_2}{\big(r_{\rm w}(s_1+\mu)\big)^2}\big(1-E(0)\big) + \frac{E(0)}{r_{\rm w}(s_1+\mu)}\big(r_{\rm w}s_2 t_N(0) + r_{\rm w}(s_1+\mu)t'_N(0)\big)\right] = 0. \tag{38}$$

Multiplying both sides by $r_{\rm w}(s_1+\mu)^2$ yields

$$-s_2(1-E_0) + E_0\,\frac{C(s_1+\mu)}{1-\mu}\,s_2 + E_0\,\frac{C}{(1-\mu)^2}(s_1+\mu)^2 = 0. \tag{39}$$

Solving above equation for $s_2$ (and evaluating the limit as $\mu \to 0$) gives *analytic separator* $s_2^\star(s_1)$ mentioned in the main text.

##### SI.1.5 Condition for substantial decrease in mutant burden with the death-to-birth ratio

To obtain a noticeable decrease in mutant burden as the death-to-birth ratio increases, we require that near $\delta \to 0^+$ the transient term has fully relaxed, i.e.,

$$t_N \gg |D|^{-1}. \tag{40}$$

Using $t_N = \ln(N/x_0)/[r_{\rm w}(1-\delta-\mu)]$ and $|D|^{-1} = 1/[r_{\rm w}|s_1+\delta s_2+\mu|]$, condition (40) becomes

$$|s_1+\delta s_2+\mu| \gg \frac{1-\delta-\mu}{\ln(N/x_0)}. \tag{41}$$

For the special case $x_0 = 1$, $s_2 = 0$, and $\mu \approx 0$, the net disadvantage of the mutant,

$$D = -r_{\rm w}(s_1+\delta s_2+\mu) \simeq -r_{\rm w}s_1,$$

remains essentially constant for all the death-to-birth ratio values. That is why the saturation condition must be satisfied already at $\delta \to 0^+$ in order to observe any decrement in the mutant burden. Under this requirement, the aforementioned condition (40) simplifies to

$$|s_1| \ll \frac{1}{\ln N}, \tag{42}$$

showing that a substantial decrease in mutant burden at small death-to-birth ratio occurs only when the selective disadvantage in division rate is much smaller than $(\ln N)^{-1}$. For another special case $x_0 = 1$, $s_1 = 0$, and $\mu \approx 0$, the net disadvantage of the mutant,

$$D = -r_{\rm w}(s_1+\delta s_2+\mu) \simeq -r_{\rm w}\delta s_2,$$

increases proportionally with the death-to-birth ratio. That is why the saturation condition (40) must be satisfied already at $\delta \to 1^-$ in order to observe a pronounced decrement in the mutant burden. To gain an estimate of how rapidly the mutant accumulation decays with increasing death-to-birth ratio, consider a intermediate representative value $\delta = 1/2$. In this case, the condition becomes

$$\left|\tfrac{1}{2}s_2\right| \gg \frac{1/2}{\ln N} \quad \Rightarrow \quad |s_2| \gg \frac{1}{\ln N}.$$

Thus, while any $s_2 < 0$ ultimately suppresses mutant accumulation at large death-to-birth ratio, a substantial and rapid decrease in mutant burden occurs only when the selective disadvantage in death rate exceeds roughly $(\ln N)^{-1}$.

##### SI.1.6 The dependence of mutant burden on $N$ and the degree of disadvantage

In Figure SI.2 we show how the dependence of $M_N$ on $\delta$ changes depending on the target population size and the degree of disadvantage. We assume that the disadvantage is in death only, that is, $s_0 = 0, s_2 < 0$. In the opposite case $s_1 < 0, s_2 = 0$, the number of mutants always increases with $\delta$. Panel (a) shows three qualitatively different regions in the $N - |s_2|$ space. Denote by $\max_\delta M_N$ and $\min_\delta M_N$ the maximum and the minimum values attained by $M_N(\delta)$ on $\delta \in [0, 1]$. For each pair $N, S_2$, in order to determine the qualitative shape of the $M_N(\delta)$ curve, we calculated the quantities $G_0 \equiv (\max_\delta M_N - M_N(0))/\min_\delta M_N$ and $G_1 \equiv (\max_\delta M_N - M_N(1))/\min_\delta M_N$. First it was determined that for all the pairs $(N, S_2)$, the difference between the minimum and the maximum value was greater than 10% of the minimum. If both $G_0$ and $G_1$ were greater than 0.1, the function $M_N$ was labeled "non-monotonic". If $G_0 < 0.1$ then it was labeled as decreasing and if $G_1 < 0.1$ it was labeled as increasing. This is what is presented in panel (a) of Figure SI.2.

Panel (b) of Figure SI.2 shows the relative change of the function $M_N(\delta)$ with $\delta$ in $\delta \in [0, 1]$. Panel (c) fixes the value of $s_2 = -10^{-1.5}$ (which corresponds to the horizontal dashed line in panel (a)), and plots the functions $M_N$ vs $N$ for several values of $\delta$, to show how the ordering changes with $N$.

#### SI.2 A stochastic, well-mixed colony.

Figure SI.1(e-h) gives a summary of the mutant burden dependency on $\delta$ for the non-spatial stochastic model, while allowing comparison with the corresponding deterministic model (panels (a-d)). Below we provide details of the methodology used to study the stochastic system.

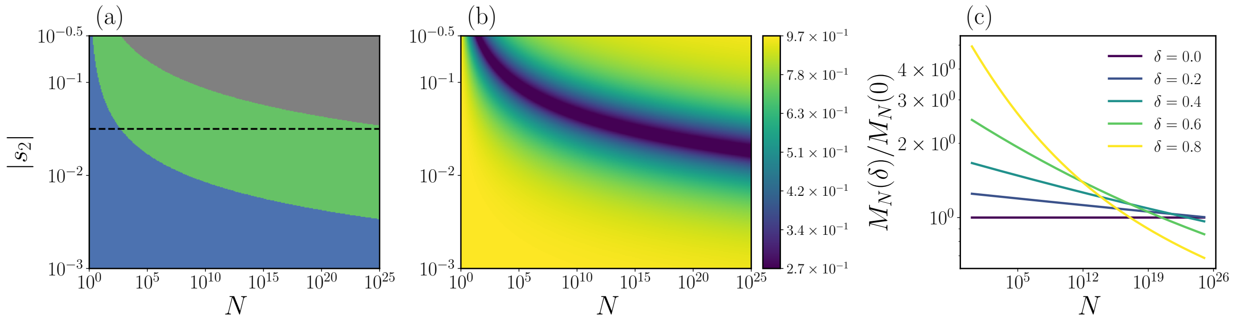

Figure SI.2: *The dependence of the deterministic model on population size and degree of disadvantage.* (a) The dependence of $M_N$ on $\delta$, as a function of $N$ and $|s_2|$. (b) The maximum relative change of $M_N$ as a function of $\delta \in [0,1]$, calculated as $(\max_\delta M_N - \min_\delta M_N)/\max_\delta M_N$. The crossover effect observed for $s_2 = -10^{-1.5}$ (corresponding to dashed horizontal line in panel (a)). Shown are the relative numbers of mutants normalized to the case $\delta = 0$, $M_N(\delta)/M_N(\delta = 0)$, for $\delta \in \{0.0, 0.2, 0.4, 0.6, 0.8\}$. Other parameters are $r_\mathrm{w} = 1$, $\mu = 10^{-7}$ and $s_1 = 0$.

##### SI.2.1 Probability of jackpot events

We study a well-mixed stochastic population that hits size $N$, consisting of wild types $X(t)$ and mutants $Y(t)$. The transition rates are

$$\begin{aligned} b_\mathrm{w} &= r_\mathrm{w}(1-\mu), \\ d_\mathrm{w} &= r_\mathrm{w}\delta, \\ b_\mathrm{m} &= r_\mathrm{w}(1+s_1), \\ d_\mathrm{m} &= r_\mathrm{w}\delta(1-s_2). \end{aligned} \tag{1}$$

The initial state is $(X(0), Y(0)) = (1, 0)$. The process reaches absorption when either $X + Y = 0$ or $X + Y = N$. Thus, there are three kinds of final outcomes: Absorption at $(X, Y) = (0, 0)$, hitting at $(X, Y) = (N - M_N, M_N)$ with $M_N < N$, or jackpots (mutant takeover) at $(X, Y) = (0, N)$. Denote these probabilities by $P^{(N)}_{0,0}$, $P^{(N)}_{X,Y}$, and the jackpot probability $P^{(N)}_{0,N}$, which satisfy

$$P^{(N)}_{0,0} + P^{(N)}_{X,Y} + P^{(N)}_{0,N} = 1. \tag{2}$$

The wild type lineage is a birth–death chain with per-individual rates $b_\mathrm{w}$ and $d_\mathrm{w}$. Define the ratio

$$R_\mathrm{w} = \frac{d_\mathrm{w}}{b_\mathrm{w}} = \frac{\delta}{1-\mu}. \tag{3}$$

For such a chain the probability that $X$ hits 0 before $N$, starting from $X = 1$ is [1]

$$q^{(N)}_X(1) = 1 - \frac{1 - R_\mathrm{w}}{1 - R_\mathrm{w}^N} = \frac{R_\mathrm{w} - R_\mathrm{w}^N}{1 - R_\mathrm{w}^N}, \tag{4}$$

Similarly, the mutant lineage has ratio

$$R_\mathrm{m} = \frac{d_\mathrm{m}}{b_\mathrm{m}} = \frac{\delta(1-s_2)}{1+s_1}. \tag{5}$$

For a single mutant, the probability of eventual extinction before reaching $N$ is

$$q_Y^{(N)}(1) = 1 - \frac{1 - R_\mathrm{m}}{1 - R_\mathrm{m}^N} = \frac{R_\mathrm{m} - R_\mathrm{m}^N}{1 - R_\mathrm{m}^N}, \tag{6}$$

and the probability of absorption at $N$ is

$$p_Y^{(N)}(1) = \frac{1 - R_\mathrm{m}}{1 - R_\mathrm{m}^N}. \tag{7}$$

While the wild type lineage is alive, each individual produces mutants at rate $\nu$. If $T_X$ is the time when the wild type lineage is absorbed at 0, then the total number of mutant individuals created up to that time is

$$M = r_\mathrm{w}\mu \int_0^{T_X} X(t)\, dt. \tag{8}$$

This integral computes the area under the curve of the wild type process, and multiplying by $\nu$ converts it into the expected number of mutation events. If the wild type lineage is *subcritical* ($b_\mathrm{w} < d_\mathrm{w}$), its mean population size decays exponentially [2] as

$$\mathbb{E}[X(t)] = e^{-(d_\mathrm{w} - b_\mathrm{w})t}. \tag{9}$$

Integrating this mean trajectory over all time gives the expected total area before extinction,

$$\mathbb{E}\left[\int_0^\infty X(t) dt\right] = \frac{1}{d_\mathrm{w} - b_\mathrm{w}}. \tag{10}$$

However, wild-type considered in our model is always supercritical. We are interested in the conditional expectation given that the wild type eventually hits 0 before $N$,

$$\Lambda^{(N)}(\delta) = \mathbb{E}[M \mid X \text{hits 0 before } N]. \tag{11}$$

In the large-$N$ limit, the wild-type process can be treated as a branching process. Its probability of extinction from a single ancestor is $R_\mathrm{w} = \delta/(1-\mu)$. Conditioning on extinction as done in [3], and comparing the transformed [4] Galton–Watson generator with the standard subcritical offspring generator yields effective rates $b_\mathrm{w}^\mathrm{eff} = b_\mathrm{w} R_\mathrm{w}$ and $d_\mathrm{w}^\mathrm{eff} = d_\mathrm{w}/R_\mathrm{w}$, so that $d_\mathrm{w}^\mathrm{eff} - b_\mathrm{w}^\mathrm{eff} = b_\mathrm{w} - d_\mathrm{w}$. Hence, for such an *effectively* subcritical process [3], the expected lifetime population size is

$$\mathbb{E}\left[\int_0^\infty X(t) dt \middle| X \text{ hits } 0\right] = \frac{1}{d_\mathrm{w}^\mathrm{eff} - b_\mathrm{w}^\mathrm{eff}}, \tag{12}$$

which after substitution gives

$$\mathbb{E}\left[\int_0^\infty X(t) dt \middle| X \text{ hits } 0\right] = \frac{1}{r_\mathrm{w}(1 - \mu - \delta)}. \tag{13}$$

Multiplying by the rate of mutation $r_\mathrm{w}\mu$ yields the mean number of seeded mutants,

$$\Lambda(\delta) = \frac{\mu}{1 - \mu - \delta}. \tag{14}$$

The number of surviving mutant lineages is then Poisson distributed with mean $p_Y^{(N)}(1)\Lambda(\delta)$. Therefore the probability that no mutant lineage survives is

$$\Pr[\text{no mutant survives}] = \exp\big(-p_Y^{(N)}(1)\Lambda(\delta)\big), \tag{15}$$

and the probability that at least one survives is

$$\Pr[\text{some mutant survives}] = 1 - \exp\big(-p_Y^{(N)}(1)\Lambda(\delta)\big). \tag{16}$$

Finally, combining with the event that the wild type hits 0 before $N$, we obtain the absorption probabilities

$$P_{0,0}^{(N)}(\delta) = q_X^{(N)}(1)\exp\big(-p_Y^{(N)}(1)\,\Lambda(\delta)\big), \tag{17}$$

$$P_{0,N}^{(N)}(\delta) = q_X^{(N)}(1)\Big(1 - \exp\big(-p_Y^{(N)}(1)\Lambda(\delta)\big)\Big), \tag{18}$$

As in our simulations we reject the extinction trajectories, the effective probability of jackpot events becomes

$$P^{\text{jackpot}}(\delta, N) = \frac{P_{0,N}^{(N)}(\delta)}{1 - P_{0,0}^{(N)}(\delta)}. \tag{19}$$

We refine the deterministic well-mixed prediction to match the outcomes of the stochastic well-mixed system as follows,

$$\langle M_N^{\text{stoch}}\rangle_{\text{theory}} = \big(1 - P^{\text{jackpot}}(\delta, N)\big)M_N^{\text{determ}} + P^{\text{jackpot}}(\delta, N)\,N. \tag{20}$$

##### SI.2.2 Evolutionary rescue with instantaneous death of all wild-type individuals at treatment onset

The scenario in which the total population reaches size $N$, consisting of both wild-type and mutant individuals, becomes significantly simpler to analyze when, at the onset of treatment, all wild-type individuals die instantaneously. This is in contrast to the previously discussed case, which incorporates the finite speed of treatment. In the present setting, we show that, without explicitly using the mutant burden distribution $\mathbb{P}(M_N = k)$ at the onset of treatment, the rescue probability can be understood solely in terms of the total number of mutants produced by the wild-type population up to the time the total population reaches the target size $N$.

In this setting, treatment acts instantaneously at the onset, eliminating all wild-type individuals, so that only pre-existing mutants present at the time the total population first reaches size $N$ can contribute to evolutionary rescue. Denoting by $M_N$ the number of mutants at that time, the rescue probability is given by

$$P_{\text{resc}}(\delta) = \sum_{k=0}^{N} \mathbb{P}(M_N = k)\, p_Y^{(\widetilde{N})}(k), \tag{21}$$

where $p_Y^{(\widetilde{N})}(k)$ is the probability that $k$ mutant individuals ultimately reaches the final size $\widetilde{N}$. As we do not have an explicit analytical expression for $\mathbb{P}(M_N = k)$, the above equation is difficult to interpret physically in terms of the behavior of rescue probabilities across different death-to-birth ratios. Hence, we adopt an alternative approach to compute the rescue probability as follows.

The mean number of mutants produced, note that this is not the mutant burden at size $N$, since mutants undergo both birth and death and hence grow dynamically, which is not required for computing mutant production by the wild-type, up to the time the total population reaches $N$ is

$$\Lambda_{\text{prod}}(\delta) = r_{\text{w}}\mu \int_0^{t_N} X(t)\, dt, \tag{22}$$

where the wild-type population grows as $X(t) \approx e^{g_{\text{w}} t}$ with growth rate $g_{\text{w}} = r(1 - \mu - \delta)$, yielding

$$\Lambda_{\text{prod}}(\delta) = \frac{r\mu}{g_{\text{w}}}(N-1) = \frac{\mu}{1-\mu-\delta}(N-1). \tag{23}$$

The rescue probability of a single mutant lineage that reaches the target size $\widetilde{N}$ is given by

$$\nu_{\text{surv}}(\delta) = \frac{1 - R_{\text{m}}}{1 - R_{\text{m}}^{\widetilde{N}}}, \tag{24}$$

where

$$R_{\text{m}} = \frac{d_{\text{m}}}{b_{\text{m}}} = \frac{\delta(1-s_2)}{1+s_1}. \tag{25}$$

The number of surviving mutant lineages is then Poisson distributed with mean $\nu_{\text{surv}}(\delta)\Lambda_{\text{prod}}(\delta)$. The probability that at least one mutant lineage rescues the population, i.e., the rescue probability, becomes

$$P_{\text{resc}}(\delta) = 1 - \exp\left(-\frac{1-R_{\text{m}}}{1-R_{\text{m}}^{\widetilde{N}}} \cdot \frac{\mu}{1-\mu-\delta}(N-1)\right) \tag{26}$$

Please note that Eqs. (21) and (26) both capture the simulation results. However, the latter more clearly reflects the underlying rescue mechanism and can be expressed explicitly in terms of the chosen parameters, making it more interpretable.

**Monotonic decrease of rescue probability with $\delta$.** We have

$$P_{\text{resc}}(\delta) = 1 - \exp\left(-\nu_{\text{surv}}(\delta)\, \Lambda_{\text{prod}}(\delta)\right). \tag{27}$$

Since $1 - e^{-x}$ is a strictly increasing function of $x$, it suffices to show that

$$K(\delta) := \nu_{\text{surv}}(\delta)\, \Lambda_{\text{prod}}(\delta) \tag{28}$$

is a decreasing function of $\delta$. Taking the derivative with respect to $\delta$,

$$\frac{dK}{d\delta} = \Lambda_{\text{prod}}(\delta)\, \frac{d\nu_{\text{surv}}(\delta)}{d\delta} + \nu_{\text{surv}}(\delta)\, \frac{d\Lambda_{\text{prod}}(\delta)}{d\delta}. \tag{29}$$

We now compute the two derivatives appearing in above equation.

$$\frac{d\Lambda_{\text{prod}}}{d\delta} = \frac{\mu(N-1)}{(1-\mu-\delta)^2} > 0. \tag{30}$$

Next, for large $\widetilde{N}$, we use the approximation $R_\mathrm{m}(\delta)^{\widetilde{N}} \to 0$, so

$$\nu_\mathrm{surv}(\delta) = \frac{1 - R_\mathrm{m}(\delta)}{1 - R_\mathrm{m}(\delta)^{\widetilde{N}}} \;\approx\; 1 - R_\mathrm{m}(\delta). \tag{31}$$

Therefore,

$$\frac{d\nu_\mathrm{surv}(\delta)}{d\delta} = -\frac{dR_\mathrm{m}(\delta)}{d\delta}. \tag{32}$$

Substituting into $dK/d\delta$,

$$\begin{aligned} \frac{dK}{d\delta} &= \Lambda_\mathrm{prod}(\delta)\left(-\frac{dR_\mathrm{m}}{d\delta}\right) + (1 - R_\mathrm{m})\frac{d\Lambda_\mathrm{prod}}{d\delta} && (33) \\ &= -\frac{\mu(N-1)}{1-\mu-\delta}\frac{dR_\mathrm{m}}{d\delta} + (1 - R_\mathrm{m})\frac{\mu(N-1)}{(1-\mu-\delta)^2}. && (34) \end{aligned}$$

Factoring,

$$\frac{dK}{d\delta} = \frac{\mu(N-1)}{(1-\mu-\delta)^2}\left[(1 - R_\mathrm{m}) - (1-\mu-\delta)\frac{dR_\mathrm{m}}{d\delta}\right]. \tag{35}$$

We now substitute $R_\mathrm{m}(\delta) = \delta(1-s_2)/(1+s_1)$ and $dR_\mathrm{m}/d\delta = (1-s_2)/(1+s_1)$ in the above equation, which gives

$$\begin{aligned} \frac{dK}{d\delta} &= \frac{\mu(N-1)}{(1-\mu-\delta)^2}\left[\left(1 - \frac{\delta(1-s_2)}{1+s_1}\right) - (1-\mu-\delta)\frac{1-s_2}{1+s_1}\right] \\ &= \frac{\mu(N-1)}{(1-\mu-\delta)^2}\left[\frac{(1+s_1) - \delta(1-s_2)}{1+s_1} - \frac{(1-\mu-\delta)(1-s_2)}{1+s_1}\right] \\ &= \frac{\mu(N-1)}{(1-\mu-\delta)^2(1+s_1)}\left[(1+s_1) - (1-\mu)(1-s_2)\right] \\ &= \frac{\mu(N-1)}{(1-\mu-\delta)^2(1+s_1)}\left[s_1 + s_2 - \mu(1-s_2)\right]. && (36) \end{aligned}$$

For a disadvantageous mutant with $(s_1 + s_2) < \mu(1 - s_2)$, we obtain $dK/d\delta < 0$, which implies

$$\frac{dP_\mathrm{resc}}{d\delta} < 0, \tag{37}$$

and the rescue probability decreases monotonically with increasing $\delta$.

##### SI.2.3 Gillespie simulation algorithm

To simulate the stochastic dynamics of the well-mixed stochastic system, we use the Gillespie algorithm, which models birth–death processes as sequences of discrete events. The reaction propensities are defined as follows:

$$\begin{aligned} a_1 &= r_\mathrm{w}(1-\mu)X, \quad \text{(wild-type reproduction)} \\ a_2 &= d_\mathrm{w}X, \quad \text{(wild-type death)} \\ a_3 &= r_\mathrm{w}\mu X, \quad \text{(mutation during division of a wild type)} \\ a_4 &= r_\mathrm{m}Y, \quad \text{(mutant reproduction)} \\ a_5 &= d_\mathrm{m}Y. \quad \text{(mutant death)} && (38) \end{aligned}$$

The algorithm proceeds as follows. The initial state $(X, Y)$ is set, together with the model parameters $r_{\mathrm{w}}, d_{\mathrm{w}}, r_{\mathrm{m}}, d_{\mathrm{m}}$, and $\mu$. The total propensity is then

$$a_0 = \sum_{i=1}^{5} a_i. \tag{39}$$

Two random numbers $r_1$ and $r_2$ are drawn independently from a uniform distribution on $(0, 1)$. The waiting time until the next event is

$$\Delta t = \frac{1}{a_0} \ln\left(\frac{1}{r_1}\right). \tag{40}$$

The next reaction is chosen by computing the cumulative sums

$$A_j = \sum_{i=1}^{j} a_i, \tag{41}$$

and selecting the smallest $j$ such that

$$A_j \geq r_2 a_0. \tag{42}$$

The system state is then updated according to the selected reaction:

$$\begin{aligned} j = 1: &\quad X \mapsto X + 1, \\ j = 2: &\quad X \mapsto X - 1, \\ j = 3: &\quad Y \mapsto Y + 1, \\ j = 4: &\quad Y \mapsto Y + 1, \\ j = 5: &\quad Y \mapsto Y - 1. \end{aligned}$$

The simulation time is updated as $t \mapsto t + \Delta t$, and the new values of $(t, X, Y)$ are recorded. This procedure is iterated until a predefined stopping condition is reached, such as the total population size hitting $X + Y = N$ or absorption at extinction of whole population. For each non-extinct trajectory, the final mutant size $M_N$ is recorded to compute the ensemble-averaged mutant burden $\langle M_N \rangle$.

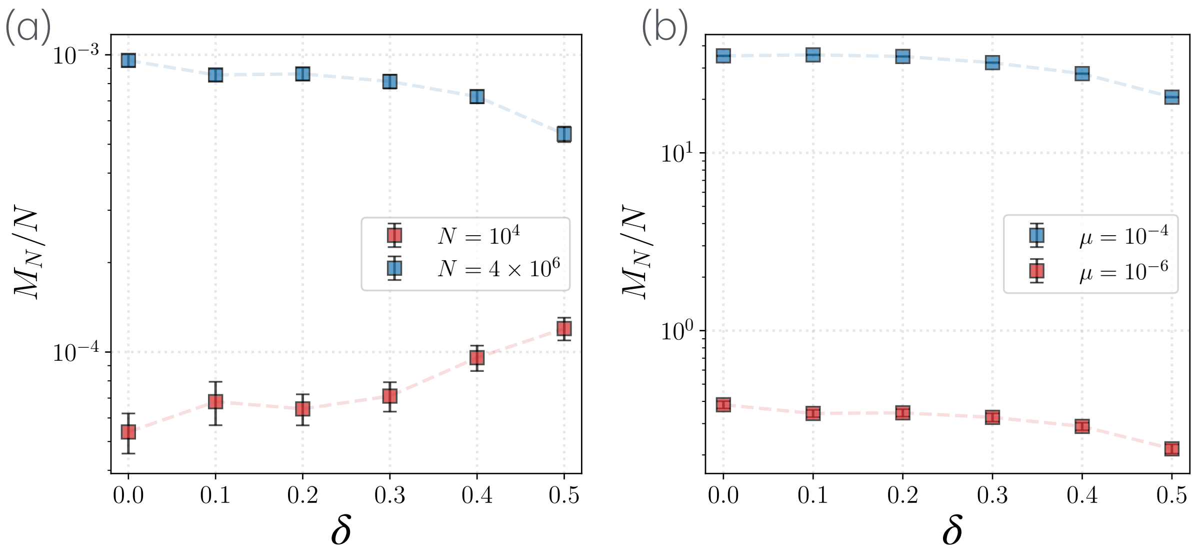

Figure SI.3: *Dependence of fractional mutant burden on the death-to-birth ratio for spatial colonies at different population sizes and mutation rates.* Panel (a): The red curve corresponds to $N = 1 \times 10^4$, and the blue curve to $N = 4 \times 10^6$; the mutation rate is $\mu = 1 \times 10^{-6}$. Panel (b): The red curve corresponds to $\mu = 10^{-6}$ and the blue curve to $\mu = 10^{-4}$; the population size is $N = 4 \times 10^6$. Vertical bars indicate the standard error. A total of $8 \times 10^4$ trajectories were simulated for $N = 1 \times 10^4$, and $1.6 \times 10^4$ trajectories were simulated for $N = 4 \times 10^6$. Other parameters are: $(s_1, s_2) = (0.00, -0.02)$, $r_{\mathrm{w}} = 0.5$.

#### SI.3 A stochastic, spatial colony

##### SI.3.1 Effects of population size

In the main text, we analyze the mutant burden for very large population sizes, small mutation rates, and mutants that are disadvantaged either in birth or in death (see Figure 3 of the main text). In both cases, we find that the mutant burden tends to decrease as the death-to-birth ratio increases. However, we now investigate what happens when the final population size is small. In this case, the dependence on the death-to-birth ratio can reverse, and the mutant burden may instead increase with increasing the death-to-birth ratio (see Figure SI.3(a)).

##### SI.3.2 Effect of Mutation Rate

Increasing the mutation rate has a straightforward effect. For any fixed death-to-birth ratio, the mutant burden increases. However, the qualitative relationship between mutant burden and the death-to-birth ratio remains unchanged. The primary effect of increasing the mutation rate is a vertical shift of the curve (see Figure SI.3(b)).

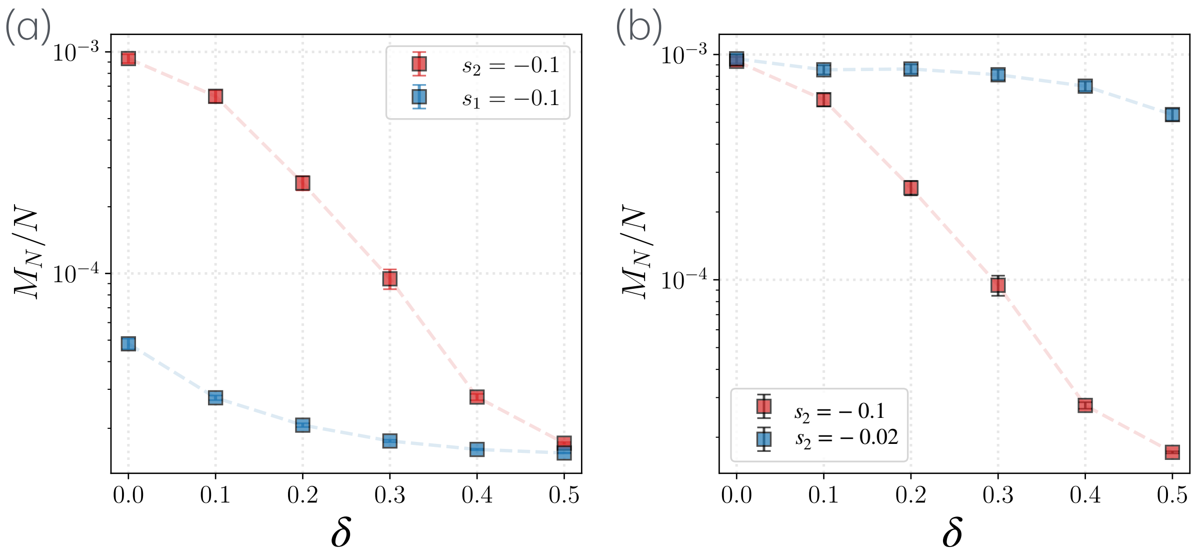

Figure SI.4: *Dependence of fractional mutant burden on the death-to-birth ratio in spatial colonies for different source and degree of disadvantage.* Panel (a): The red curve corresponds to disadvantage parameters $(s_1, s_2) = (-0.10, 0.00)$, and the blue curve to $(s_1, s_2) = (0.00, -0.10)$. Panel (b): The red curve corresponds to disadvantage parameters $(s_1, s_2) = (0.00, -0.10)$, and the blue curve to $(s_1, s_2) = (0.00, -0.02)$. Parameters are $\mu = 10^{-6}$ and $r_{\mathrm{w}} = 0.5$. Vertical bars represent the standard error. The population size is $N = 4 \times 10^6$, and a total of $1.6 \times 10^4$ independent trajectories were simulated.

#### SI.3.3 Effects of selection strength

We also compared two types of selection of disadvantageous mutants in isolation: disadvantage in birth and disadvantage in death. We find that when the disadvantage acts only through death, the mutant burden decreases much more sharply with increasing the death-to-birth ratio compared to the case where the mutant is disadvantaged only in birth (see Figure SI.4(a)). We also compared the effect of the strength of disadvantage when mutants are disadvantaged in death. We observed that a stronger death disadvantage leads to a lower mutant burden at high death-to-birth ratio (see Figure SI.4(b)). One can similarly verify that this qualitative behavior holds when mutants are disadvantaged only in birth.

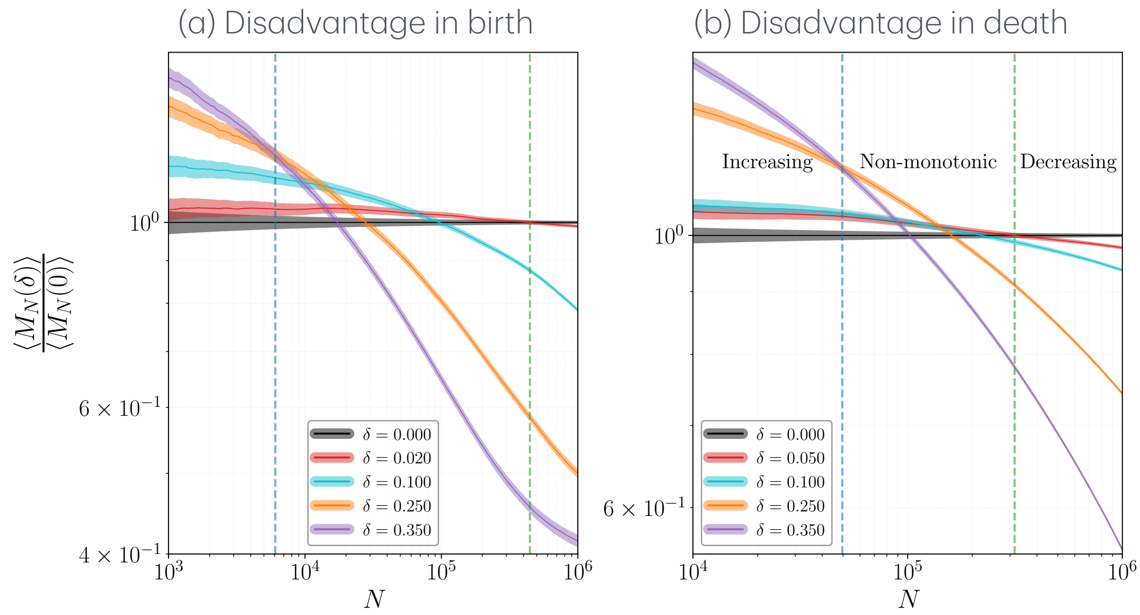

Figure SI.5: *Crossovers of disadvantageous mutant burden (normalized by the mutant burden at the zero death-to-birth ratio) for different death-to-birth ratios as the final population size increases.* (a) The normalized mutant burden as a function of final population size $N$ for a mutant with disadvantage acting only through birth, $(s_1, s_2) = (-0.10, 0.00)$. (b) Same as (a), but for a mutant with disadvantage acting only through death, $(s_1, s_2) = (0.00, -0.05)$, highlighting the crossover behavior between different death-to-birth ratios. In (a–b), different colors represent different death-to-birth ratios. Each curve represents the average over $10^5$ independent simulations. Solid lines represent the mean, and the transparent thickness represents the standard error. Other parameters are $r_{\mathrm{w}} = 0.5$ and $\mu = 10^{-4}$.

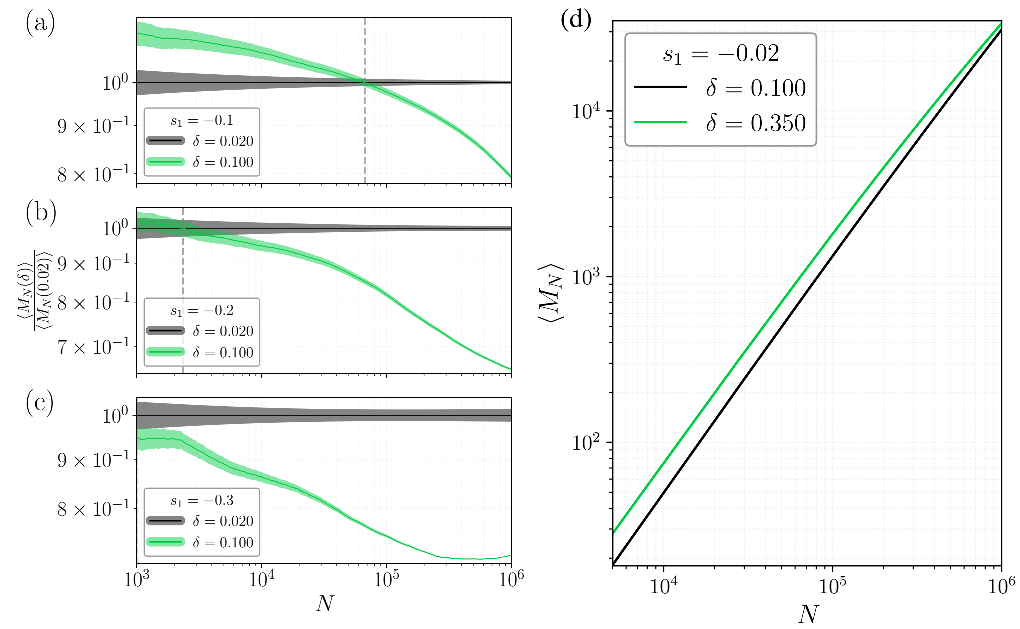

Figure SI.6: *Crossovers of disadvantageous mutant burden for different death-to-birth ratios as the final population size increases.* (a) The mutant burden (normalized by the mutant burden at the death-to-birth ratio $\delta = 0.02$) as a function of final population size $N$ for a mutant with disadvantage acting only through birth, $(s_1, s_2) = (-0.10, 0.00)$. (b) Same as (a) but with a stronger disadvantage, $(s_1, s_2) = (-0.20, 0.00)$. (c) Same as (a–b) but with an even stronger disadvantage, $(s_1, s_2) = (-0.30, 0.00)$. In (a–c), different colors represent different death-to-birth ratios. As the disadvantage increases, the crossover between different death-to-birth ratios (vertical dashed line) occurs at progressively smaller final population sizes. Each curve represents the average over $10^5$ independent simulations. Solid lines represent the mean, and the transparent thickness represents the standard error. In panel (d), the mutant burden is shown for two different death-to-birth ratios for mutants with a small disadvantage, $(s_1, s_2) = (-0.02, 0.00)$. Because the disadvantage is weak, no clear crossover in mutant burden is observed as the population size is increased up to $10^6$. Other parameters are $r_{\mathrm{w}} = 0.5$ and $\mu = 10^{-4}$.

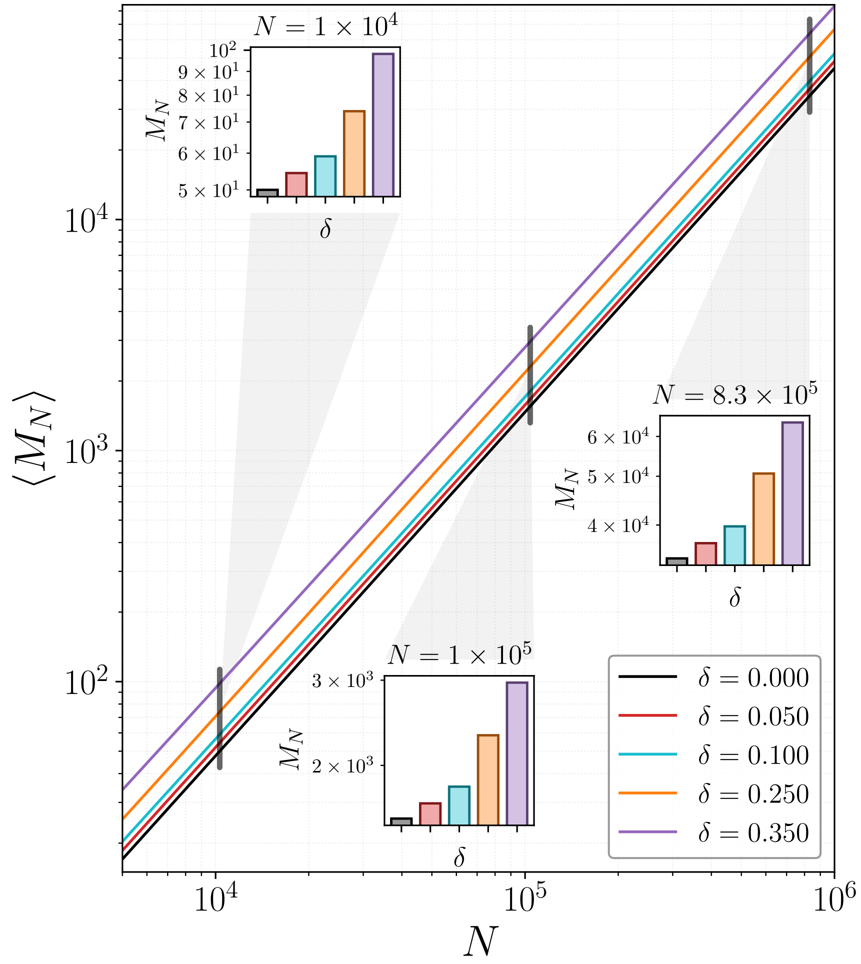

Figure SI.7: *Dependence of the neutral mutant burden on the final population size.* The expected number of mutants, $\langle M_N \rangle$, is shown as a function of the final population size $N$ for different death-to-birth ratios $\delta$, indicated by different colors. Each curve represents the average over $10^5$ independent simulations. Other parameters are fixed at $r_{\mathrm{w}} = 0.5$ and $\mu = 10^{-4}$.

##### SI.3.4 Mutant burden cross-overs

To illustrate the mutant burden crossovers corresponding to different death-to-birth ratios discussed in the main text, we normalize the mutant burden at each death-to-birth ratio value by the mean burden observed in the zero-death-to-birth ratio case. This normalization allows us to clearly visualize the crossovers and how the dependence of mutant burden on the death-to-birth ratio changes as the final population size increases. In Figure SI.5, three distinct regimes can be identified. For small final population sizes, the mutant burden increases monotonically with the death-to-birth ratio. For intermediate population sizes, a non-monotonic dependence on the death-to-birth ratio emerges. Finally, for large final population sizes, the trend reverses, and the mutant burden decreases monotonically with the death-to-birth ratio. An increase in the strength of the disadvantage causes the crossover to occur at smaller final population sizes (see Figure SI.6(a-c)). For a very small disadvantage, the crossover occurs only at very large population sizes; see Figure SI.6(d). We also observe no crossover for neutral mutants (see Figure SI.7).

##### SI.3.5 Evolutionary rescue and non-viability of mutants in the spatial model

In the rescue framework, mutant viability plays a central role. Increasing the death-to-birth ratio raises the effective death rate experienced by mutants, thereby enhancing lineage extinction and eventually rendering mutants non-viable. Once mutants become non-viable, the probability of rescue must vanish. In the well-mixed model with a twenty percent disadvantage in birth, mutants lose viability at $\delta = 0.8$. In the spatial model, however, additional constraints arise from spatial exclusion: Birth events are limited by local crowding in the bulk, effectively reducing the reproductive potential of mutants. As a result, mutants are expected to become non-viable at smaller death-to-birth ratio values than in the well-mixed case.

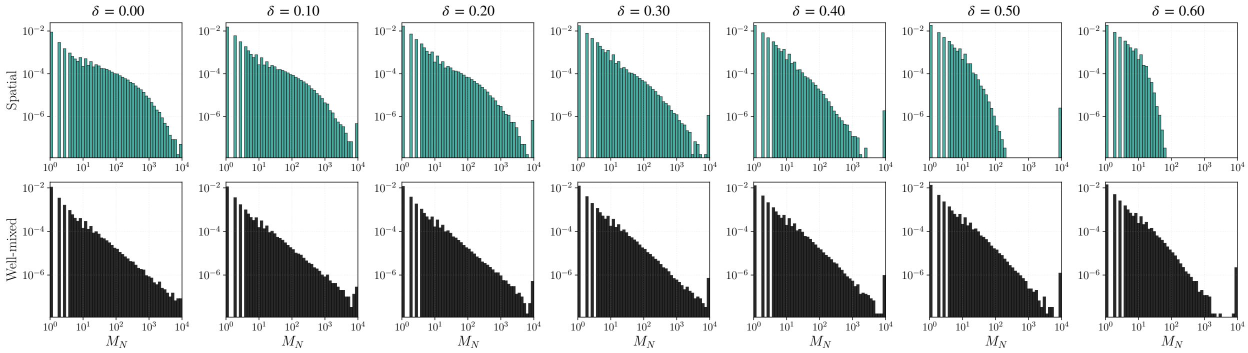

Figure SI.8: *Probability distribution of mutant numbers at threshold size $N$.* We consider mutants that have selective disadvantage only in birth ($s_1 = 0.20$), while having no disadvantage in death ($s_2 = 0$). The final population size is fixed at $N = 10^4$. The mutation rate is $\mu = 2 \times 10^{-6}$ and the wildtype birth rate is $r_{\mathrm{w}} = 0.25$. The probability of observing zero mutants is excluded from this plot. Each histogram is obtained from $2 \times 10^6$ independent simulations that remained non-extinct up to the threshold size. The top row represents outcomes for the spatial system, and the bottom row shows the corresponding results for the well-mixed colony at different death-to-birth ratio values.

This expectation is supported by the histograms in Figure SI.8. At $\delta = 0.6$, no takeover events by jackpot mutants are observed in the spatial system; specifically, at the threshold size there are no realizations in which the population consists entirely of mutants. In the well-mixed stochastic model, such takeover events can occur only when mutants are viable. By analogy, the absence of takeover events in the spatial model provides a practical indicator of loss of viability. Consequently, at the largest death-to-birth ratio value considered, mutants are effectively non-viable in the spatial system, and the rescue probability becomes exactly zero.
